# Improving bioplastic production by *Rhodopseudomonas palustris* TIE-1 using synthetic biology and metabolic engineering

**DOI:** 10.1101/2023.05.17.541174

**Authors:** Tahina Onina Ranaivoarisoa, Wei Bai, Karthikeyan Rengasamy, Hope Steele, Miriam Silberman, Jennifer Olabode, Arpita Bose

## Abstract

With the increasing demand for sustainably produced renewable resources, it is important to look towards microorganisms capable of producing bioproducts such as biofuels and bioplastics. Though many systems for bioproduct production are well documented and tested in model organisms, it is essential to look beyond to non-model organisms to expand the field and take advantage of metabolically versatile strains. This investigation centers on *Rhodopseudomonas palustris* TIE-1, a purple, non-sulfur autotrophic, and anaerobic bacterium capable of producing bioproducts that are comparable to their petroleum-based counterparts. To induce bioplastic overproduction, genes that might have a potential role in the PHB biosynthesis such as the regulator, *phaR,* and *phaZ* known for its ability to degrade PHB granules were deleted using markerless deletion. Mutants in pathways that might compete with polyhydroxybutyrate (PHB) production such as glycogen and nitrogen fixation previously created to increase *n*-butanol production by TIE-1 were also tested. In addition, a phage integration system was developed to insert RuBisCO (RuBisCO form I and II genes) driven by a constitutive promoter *P_aphII_* into TIE- 1 genome. Our results show that deletion of the *phaR* gene of the PHB pathway increases PHB productivity when TIE-1 was grown photoheterotrophically with butyrate and ammonium chloride (NH_4_Cl). Mutants unable to make glycogen or fix dinitrogen gas show an increase in PHB productivity under photoautotrophic growth conditions with hydrogen. In addition, the engineered TIE-1 overexpressing RuBisCO form I and form II produces significantly more polyhydroxybutyrate than the wild type under photoheterotrophy with butyrate and photoautotrophy with hydrogen. Inserting RuBisCO genes into TIE-1 genome is a more effective strategy than deleting competitive pathways to increase PHB production in TIE-1. The phage integration system developed for TIE-1 thus creates numerous opportunities for synthetic biology in TIE-1.

## Introduction

Recent improvements in genetic engineering tools have enabled scientists to systematically engineer organisms that produce various value-added chemicals, including biofuels, therapeutic products, food, and bioplastics [1–3]. In the early years, most of these engineering efforts were focused on widely used model organisms, such as *Escherichia coli*, *Saccharomyces cerevisiae,* and *Synechococcus* sp. [4–8]. This led to the development of a wide range of genetic tools for these organisms, which have been used in physiology studies and the production of useful biomolecules [9–14]. However, these heterotrophic model organisms can mostly use organic carbon as carbon source, leading to high bioproduction costs [15–17]. Many engineering efforts have used these model organisms to synthesize valuable compounds and increase their bioproduction [8, 18–22]. Recent studies have shown numerous advantages of using other microbial chassis besides model organisms for bioproduction [23–25]. Thus, in recent years more studies have focused on expanding host choice for synthetic biology, moving toward non-model organisms. One of the microbes that has drawn attention during the past decade is *Rhodopseudomonas palustris* TIE-1 (TIE-1) [26] [27, 28]. TIE-1 is a gram-negative purple non-sulfur photosynthetic bacterium known for its versatile metabolism, making it a great host for different bioproduction and pathway studies [26, 27]. TIE-1 has four primary metabolisms: chemoautotrophy, photoautotrophy, chemoheterotrophy, and photoheterotrophy [26, 27]. These different metabolisms enable TIE-1 to use a wide variety of carbon sources such as carbon dioxide (CO_2_) and many organic acids. TIE- 1 also can use ammonium salt or fix nitrogen from dinitrogen gas (N_2_) as a nitrogen source [26] . Moreover, multiple electron sources including hydrogen (H_2_), ferrous iron Fe(II), and poised electrodes are used by TIE-1 [26, 29]. One of the most appealing features of TIE-1 is its ability to uptake electrons directly from a poised electrode, which enables us to combine synthetic biology with microbial electrosynthesis (MES) [27, 30–32]. MES is a system in which microorganisms are used to produce valuable compound using their ability to exchange electrons from the bioelectrical reactor [30, 32, 33]. Using electrons from a poised electrode or a solar panel-powered MES system, TIE-1 produced biodegradable plastic and biofuel using CO_2_ as a carbon source, N_2_ as a nitrogen source, and light as an energy source [34]. This represented the first step toward a sustainable and carbon-neutral process for producing biofuel via MES using TIE-1. Besides its ability to utilize various substrates for bioproduction, TIE-1’s metabolic diversity also makes it an extraordinary model organism for pathway investigation [35, 36]. For example, the use of RuBisCO mutants allowed us to study the link between the Calvin-Benson-Bassham (CBB) cycle in carbon fixation and the extracellular electron transport[37]. Similarly, a TIE-1 *pioABC* mutant was used to investigate the electron uptake mechanism during photoferrotrophy and electrotrophy [35–37]. Not only do these studies provide insight into TIE-1’s metabolism, but they also open doors for a deeper understanding of other closely related purple non-sulfur bacteria, such as CGA009 that has been studied extensively for biohydrogen production [38, 39]. However, compared to the model organisms cited earlier, the number of genetic tools for TIE-1 is limited to those utilizing homologous recombination [31, 37].

Homologous recombination results in markerless deletion which is one of its biggest advantages. **Figure S1** shows the two steps during gene deletion using this technique. To produce *n*-butanol in TIE-1, a nitrogen fixation mutant lacking two copies of the transcriptional activators *nifA* (Rpal_1624 & Rpal_5113) has been previously generated using the markerless deletion technique. The *nifA* double mutant carrying the five gene cassette for *n*-butanol biosynthesis showed higher butanol production compared to the wild type [34]. Besides *n*-butanol, we also have reported the ability of the wild type TIE-1 and two *Rhodomicrobium* species to produce bioplastic polyhydroxybutyrate (PHB) [21][Conners et al, 2023, Manuscript in preparation]. Due to the plastic crisis in recent years, bioplastics have become an active area of research (reviewed in [40]). PHB is a member of the well-studied polyhydroxyalkanoate (PHA) family. They preserve the advantages of petroleum-based plastics, such as high durability, moldability, water, and heat resistance while being biocompatible [41–43]. Furthermore, many microorganisms can naturally degrade bioplastics within 5 – 6 weeks, while petroleum-based plastics can take hundreds to thousands of years to degrade [42–44]. However, the high feedstock cost remains an obstacle to bioplastic’s competitiveness in the market (reviewed in [45, 46]). This issue can be addressed by using photoautotrophic microbes that can use cheap alternative and waste feedstock (such as CO_2_) to produce PHA [27].

The PHB pathway consists mainly of *phaA, phaB, phaC, phaZ* genes*. phaA* encodes a ß-keto thiolase while *phaB* encodes an acetoacetyl CoA reductase. PHB polymerization is performed by PhaC whereas its mobilization during carbon demand is performed by the PHB depolymerase, PhaZ. Additional proteins have also been reported to contribute to the maturation and regulation of PHB granules by phasin (PhaP) and (PhaR), respectively [20, 27]. We have previously reported that TIE-1 possesses one *phaR* gene: Rpal_0531 using bioinformatics [27]. In *Paracoccus denitrificans* PhaR is a repressor of PHB synthesis, and acts by binding to the intergenic region of the *phaC-phaP* and *phaP-phaR* genes [47]. However, PhaR is proposed to be an activator for PHB synthesis in *Cupriavidus necator* in a PhaP-dependent and independent manner. Deletion of the PhaR gene decreased PHB production in the organism [48]. We use a deletion method here to investigate the role of PhaR in PHB biosynthesis in TIE-1.

To improve PHB production, several gene manipulations have been undertaken on the genes directly involved in the PHB pathway or genes from other pathways that may impact or compete with the PHB biosynthesis pathway [49–51]. For instance, in one study, *Sinorhizobium meliloti* was shown to produce PHB using formate generated via electrochemical CO_2_ reduction [49]. The authors then showed that PHB production increased following the deletion of the PHB depolymerase, *phaZ,* compared to the wild type [49]. This suggests that prevention of PHB degradation results in intracellular accumulation of PHB in the *phaZ* mutant. Metabolic pathways such as glycogen production and nitrogen fixation are also potential competitors for bacterial PHB production. In a monoculture of the model organism *Synechocystis* sp. PCC 6803, PHB accumulation is reported to be linked to glycogen production under prolonged nitrogen starvation [50]. Mutants lacking the glycogen phosphorylase genes showed impaired PHB accumulation, supporting the link between glycogen and PHB synthesis under nitrogen starvation (NaNO_3_) conditions in PCC 6803 [50]. Nitrogen fixation is a highly demanding electron pathway. PHB accumulation has also been reported to be triggered by nitrogen-deprived ((N_2_) or NH_4_Cl) in many bacteria [38, 52, 53]. In a parallel effort to increase PHB production, carbon fixation through the Calvin-Benson-Bassham (CBB) cycle in *Ralstonia eutropha* (now *C. necator*) has been enhanced. The RuBisCO gene from *Synechococcus* sp. PCC 7002 was overexpressed heterologously. This increased cell density (measured by optical density) by up to 89.2% and augmented the mass percent of PHB production by up to 99.7% [54].

To expand the genetic engineering tools used in TIE-1, we explored phage recombination techniques to integrate relevant genes into its genome for enhancing PHB production. Phage recombination has gained attention for genome engineering in various bacteria, including *Methanosarcina* [55] and *Mycobacterium smegmatis* [56], due to its simple design and high efficiency [57, 58]. Among different phage recombinases, φC31 has been frequently used because it does not require a helper protein and the recombination is unidirectional [59]. This system is widely used in genome engineering. For example: in *Clostridium ljungdahlii*, a whole butyric acid synthesis pathway was integrated into its genome by φC31 recombinase [57]; in *Methanosarcina* spp. the φC31 recombinase reached genome editing efficiency that is 30 times higher than homologous recombination [55].

We used the φC31 integration system we developed here to integrate additional copies of the two RuBisCO genes form I and II driven by *P_aphII_* constitutive promoter into its genome. Additionally, to increase PHB accumulation in TIE-1, we designed mutants lacking genes in the PHB pathway, i.e., the PHB depolymerase *phaZ* and the PHB regulator *phaR* genes. Δ*phaR* also helped us elucidate whether *phaR* is potentially an activator or repressor of the PHB pathway in TIE-1. Mutants lacking genes in pathways hypothesized to be competitive to the PHB synthesis pathway that were generated previously were also tested for PHB production. These include the glycogen pathway (Δ*gly* mutant lacking glycogen synthase) and the nitrogen fixation pathway (*nifA*; Rpal_1624 & Rpal_5113 mutant lacking the two *NifA* regulators) [34]. We also hypothesize that like *R. eutropha,* overexpressing RuBisCO genes (form I and II) in TIE-1 by integrating them into its genome using a phage integration system could increase intracellular carbon abundance, and hence increase PHB accumulation. We test PHB production in all the strains under a variety of growth conditions. We also grow the strains under non-nitrogen fixing conditions with NH_4_Cl (referred to as growth with NH_4_Cl throughout) or under nitrogen-fixing conditions with N_2_ gas (referred to as nitrogen-fixing conditions throughout).

Our results show that deletion of *phaR* gene increased PHB productivity when TIE-1 was grown photoheterotrophically with butyrate and NH_4_Cl. We also observed an increase in PHB productivity from the glycogen and *nifA* mutants under photoautotrophic growth conditions with H_2_ and NH_4_Cl. PHB productivity increased in engineered TIE-1 strains overexpressing RuBisCO genes form I and form II under photoheterotrophy with butyrate and photoautotrophy with H_2_, irrespective of the nitrogen source used. We show that both gene deletion and overexpression can enhance PHB production by TIE-1. Moreover, our findings from TIE-1 also inform genetic engineering to improve bioplastic production in other photosynthetic bacteria in the future.

## Results

### Generating TIE-1 mutants and suicide plasmid with the different antibiotic marker

To improve polyhydroxybutyrate (PHB) production, we constructed the following mutant strains using the suicide plasmid *pJQ200KS* carrying a gentamycin antibiotic resistance cassette: 1) a mutant lacking the regulator of the PHB pathway (Δ*phaR*), and 2) a mutant lacking the PHB depolymerase (Δ*phaZ* mutant) (**Figure S1&S2**). We also tested the previously constructed TIE-1 mutant lacking the glycogen gene (Δ*gly*), as well as the double mutant lacking the nitrogen-fixing regulator *nifA* gene (Δ*nifA*) . These strains are listed in **Table 1** and the plasmids used to create them are listed in **Table 2**. We also expanded the library of our suicide plasmid vector by creating two additional suicide plasmid vectors using *pJQ200KS* as a backbone. TIE-1 has a very high resistance to gentamycin with a minimum inhibitory concentration (MIC) of 200μg/L. To make PHB production more competitive on a large scale, this high antibiotic use can be addressed by using other cheaper antibiotics, to which TIE-1 has a lower MIC. Accordingly, we constructed suicide plasmids containing two widely used antibiotic resistance markers, chloramphenicol (MIC of 100μg/L) or kanamycin (MIC of 50μg/L), using *pJQ200KS* as the backbone as shown in **Figure S3**.

**Table 1.** Strains used in this study.

| Strains | Relative characteristics | Source |
| --- | --- | --- |
| AB415 | Wild type *Rhodopseudomonas palustris* TIE-1 | [26] |
| WB065 | Wild type TIE-1 with *attB* | This study |
| WB068 | Wild type TIE-1 with $\phi C31$ integrase; *P<sub>lac</sub>*; *lacIq*; *attB* | This study |
| WB090 | Wild type TIE-1 with *mCherry*; *P<sub>aphII</sub>* promoter; *fd* terminator; *attL*; *attR* | This study |
| WB091 | Wild type TIE-1 with *loxP-mCherry-loxP*; *P<sub>aphII</sub>* promoter; *fd* terminator; *attL*; *attR* | This study |
| AB186 | TIE-1 ΔRpal_0531 (*phaR* mutant ) | This study |
| AB190 | TIE-1 ΔRpal_0578 (*phaZ* mutant ) | This study |
| WB092 | Wild type TIE-1 with RuBisCO form I overexpressed | This Study |
| WB095 | Wild type TIE-1 with RuBisCO form I and II overexpressed | This Study |
| AB188 | TIE-1 Δ*gly* mutant | [34],This study |
| AB187 | TIE-1 Δ*nifA* double mutant | [34],This study |

**Table 2.** Plasmids used in this study.

| Plasmid | Relative characteristics | Source |
| --- | --- | --- |
| pJQ200KS | ori *P15A*; Gen<sup>r</sup> | [60] |
| pAB314 | pJQ200KS with *GlmUSX*_UP and *GlmUSX*_DN | [31] |
| pAB356 | *Cm*<sup>r</sup> | [61] |
| pAB357 | *Kan*<sup>r</sup> | [61] |
| pAB358 | *Tc*<sup>r</sup> | [61] |
| pAB359 | *Amp*<sup>r</sup> | [61] |
| pSRKGm | *P<sub>lac</sub>*; *lacIq*; ori *pBBR1*; Gen<sup>r</sup> | [61] |
| pWB083 | pAB314 with *attB* | This study |
| pWB081 | pJQ200KS with *mCherry*; *P<sub>aphII</sub>* promoter; *fd* terminator and *attP* | This study |
| pWB084 | pSRKGm with *φC31* integrase | This study |
| pWB086 | pAB314 with *φC31* integrase; *P<sub>lac</sub>*; *lacIq*; *attB* | This study |
| pWB088 | pJQ200KS with *φC31* integrase; *P<sub>aphII</sub>* promoter; *fd* terminator and *attP* | This study |
| pWB091 | pJQ200KS with Kan<sup>r</sup> | This study |
| pWB092 | pJQ200kKS with Cm<sup>r</sup> | This study |
| pWB107 | Rubisco form I and form II overexpression | This study |
| pWB108 | Rubisco form I overexpression | This study |
| pAB863 | pJQ200KS with *phaR* 1kb up and 1kb down for deletion of *phaR* | This study |
| pAB868 | pJQ200KS with *phaR* 1kb up and 1kb down for deletion of *phaZ* | This study |

### Engineering TIE-1 using a phage integration system to increase PHB production in TIE-1

Because homologous recombination is the only genetic tool available for TIE-1, we created a phage integration system to integrate genes into its genome. As shown in **Figure 1**, a φC31 recombinase system requires three parts: *attB* site, *attP* site, and the φC31 integrase.

**Figure 1.**
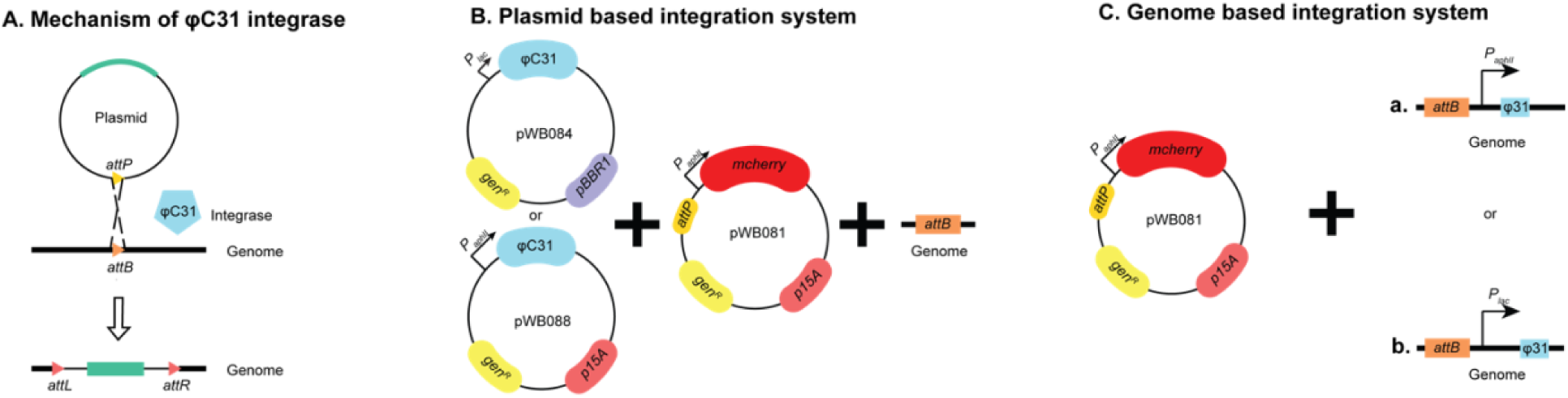
Phage integration system **A.** φC31 integrase mechanism **B.** Plasmid based integration system **C.** Genome-based integration system. *P_lac_*: *P_lac_* promoter, *_PaphII_*: *P_aphII_*promoter, *mcherry*: red fluorescent protein, *gen^R^*: gentamicin resistance, *p15A*: the origin of replication, *pBBR1*: broad host origin of replication *attP*: *attP* site for φC31integrase, *attB*: *attB* site for φC31 integrase, φC31: φC31 integrase.

Because TIE-1 does not have an *attB* site, an *attB* site was introduced into its genome. The *attP* site was introduced into a suicide plasmid with a constitutively expressed *mCherry* gene under the *P_aphII_* promoter (pWB081). For the expression of φC31 integrase, the easiest way would be to use a temperature-sensitive plasmid [62]. Unfortunately, there is no known temperature-sensitive plasmid that replicates in TIE-1. Hence, we decided to build two different systems: (1) a plasmid-based system, where the integrase is introduced into TIE-1 by a plasmid (**Figure 1B**), and (2) a genome-based system, where the integrase is integrated into the TIE-1 genome (**Figure 1C**). The advantage of the plasmid-based system is its mobility, while the genome-based system is more stable and does not rely on antibiotics. For the promoter choice, we tried both an inducible promoter (*P_lac_*) and a strong constitutive promoter (*P_aphII_*). To summarize, as shown in **Figure 1**, we have four different designs for expression of the φC31 integrase: a) *P_aphII_*-driven φC31 integrase on a suicide plasmid (pWB088); b) *P_lac_*-driven φC31 integrase on a self-replicating plasmid (pWB084); c) *P_aphII_*-driven φC31 integrase on TIE-1 genome (**Figure 1C(a)**; and d) *P_lac_*-driven φC31 integrase on TIE-1 genome (**Figure 1C(b)**).

After three separate trials, we were not able to obtain a genome-based system using the constitutive promoter (**Figure 1C(a)**), which indicated that the strong constitutive expression of φC31 integrase is probably lethal to TIE-1. Thus, we only present the results of the other three systems. As shown in **Figure 1**, the successful integration of *mCherry* was indicated by visualizing red fluorescence. All the colonies that show red fluorescence signals were tested by a PCR and 100% of these colonies showed the expected band. After the confirmation of integration, the efficiency of the different systems was evaluated by the transformation efficiency (detailed calculation described in the **Materials and Methods** section) and the integration efficiency, defined as the percentage of colonies that have red fluorescence signals among all obtained colonies.

As shown in **Figure 2A** and **2B**, the transformation efficiency normalized to the plasmid concentration is higher for the genome-based system (**Figure 1C**). This higher efficiency could be due to the sufficiency of only one plasmid for the system to be functional. Between the two plasmid-based systems, the constitutively expressed φC31 reached higher efficiency (**Figure 1B**, pWB088). This higher efficiency could be due to the independence from using the inducer, IPTG. As for the integrating frequency, the genome-based system (**Figure 1C**) and the plasmid-based system with the constitutive promoter (**Figure 1B**, pWB088) resulted in editing efficiencies of 100% (**Figure 2B**). However, the plasmid-based system with the inducible promoter only reached an editing efficiency of about 80% (**Figure. 2B**). To sum up, the genome-based system and constitutive promoter result in higher electroporation efficiency and genome editing efficiency.

After successfully integrating *mCherry*, we wanted to use the phage integration tool to assist in improving PHB production in TIE-1. Previous research conducted in *Ralstonia eutropha* (now *C. necator*) has shown that overexpression of RuBisCO resulted in higher PHB production [54]. Thus, to improve PHB production, we integrated a *P_aphII_ -*driven RuBisCO form I alone or RuBisCO form I and form II together into the TIE-1 genome to obtain two new TIE-1 strains: *Ωrub(I)* and *Ωrub(I&II)*. We were unable to obtain a plasmid with *P_aphII_ -*driven RuBisCO form II. The successful integration of these genes and *P_aphII_* were checked by PCR amplifying the *P_aphII_* region and the RuBisCO gene form I and II as indicated in **Figure S4**.

### Deletion of *phaR, phaZ*, or overexpression of the RuBisCO form I impaired the growth of TIE-1 during photoheterotrophic growth with butyrate

Because TIE-1 has previously shown high PHB production under anoxic photoheterotrophic conditions with butyrate [27], we tested this growth condition using all the constructed strains, i.e., both deletion mutants and integrants overexpressing genes of choice. Under these anoxic growth conditions, light is used as an energy source and butyrate is used both as electron and carbon sources. For all growth conditions tested, we determined the effect of nitrogen fixation by supplying N_2_ gas in the headspace (nitrogen-fixing) or ammonium chloride (NH_4_Cl) salt in the liquid media (non-nitrogen-fixing). Our results show that deleting *phaR* or *phaZ* delayed the generation time of TIE-1 by around 3 hours compared to the wild type when grown photoheterotrophically with butyrate in the presence of NH_4_Cl (**Table 3 and** **Figure 3A**). However, a slightly shorter generation time (3 hours shorter than the wild type) was obtained from the glycogen and the *nifA* double mutants. Under nitrogen-fixing growth conditions, the Δ*phaZ* mutant continued to show a growth defect observed as a longer generation time (3 hours longer than the wild type), whereas the Δ*phaR* and Δ*gly* mutants showed the same generation time as the wild type. As expected, the double mutant strain lacking the two *nifA* genes was not able to grow under nitrogen-fixing conditions (**Table 3 and** **Figure 3B**). We then compared the growth of the engineered RuBisCO strains Ω*rub(I)* and Ω*rub(I&II)* under the same photoheterotrophic growth condition with butyrate. The engineered strains carrying only the overexpressed RuBisCO gene form I (Ω*rubI*) appear to have a growth defect under both NH_4_Cl and N_2_ (**Figure 3C** **& D**). This defect seems to be more noticeable under nitrogen-fixing growth. The Ω*rub(I)* strains have an almost two-fold longer lag time than the wild type when grown photoheterotrophically with butyrate and NH_4_Cl, and almost five times longer generation time under nitrogen-fixing conditions (**Table 3, and** **Figure 3C** **& 3D**).

**Figure 3.**
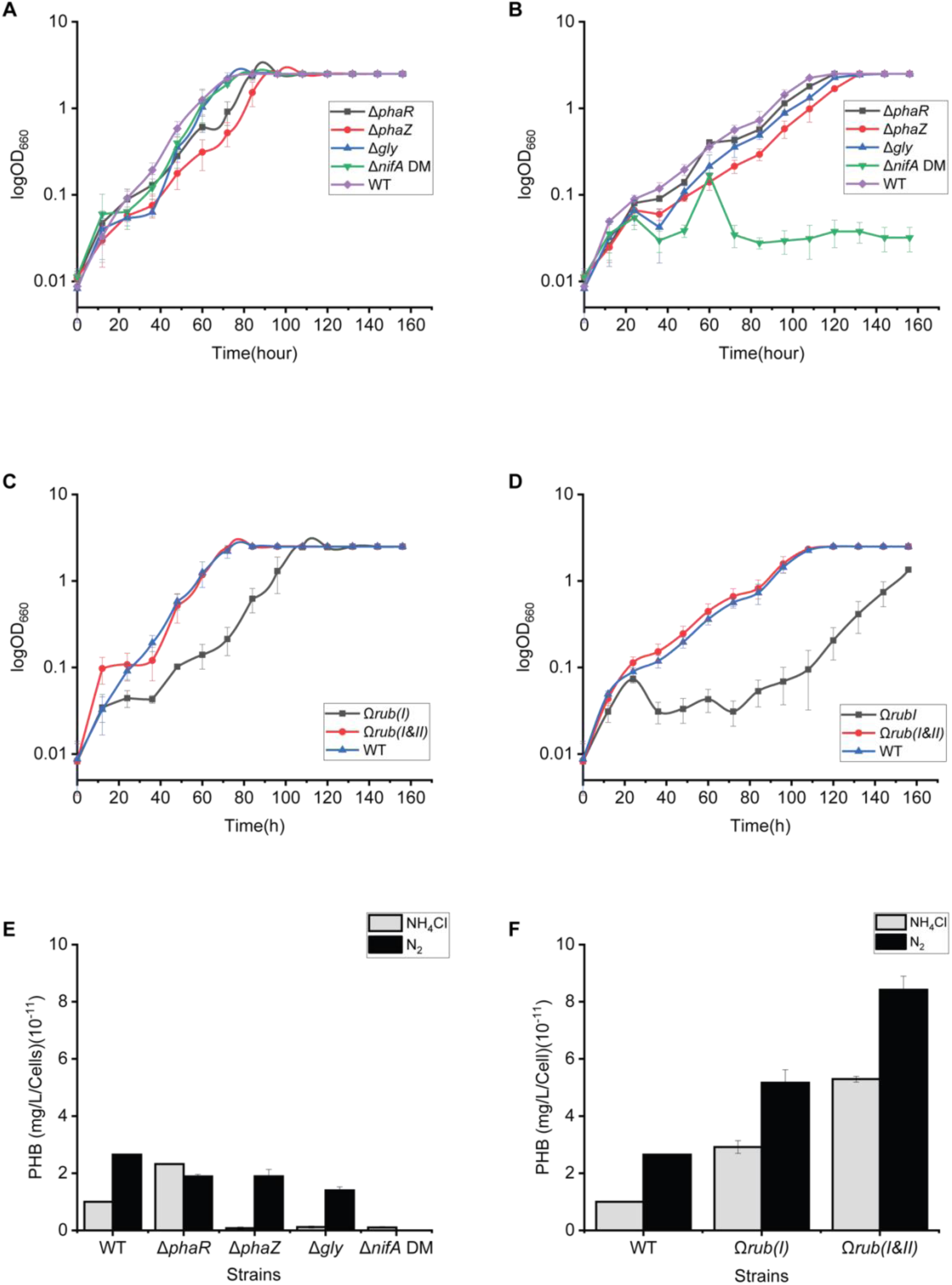
Growth and PHB productivity from different strains grown photoheterotrophically in fresh water basal media with butyrate. **A.** Growth of all mutants with NH_4_Cl. **B.** Growth of all mutants under - conditions (N_2_). **C.** Growth of the RuBisCO engineered strains with NH_4_Cl. **D.** Growth of all the RuBisCO engineered stains under nitrogen-fixing conditions. **E.** PHB productivity from mutants and TIE strains grown with butyrate with NH_4_Cl or N_2_. **F.** PHB productivity from the RuBisCO engineered and wild type TIE-1 strains grown with butyrate and NH_4_Cl or N_2_. Data are from biological triplicates.

**Table 3.**
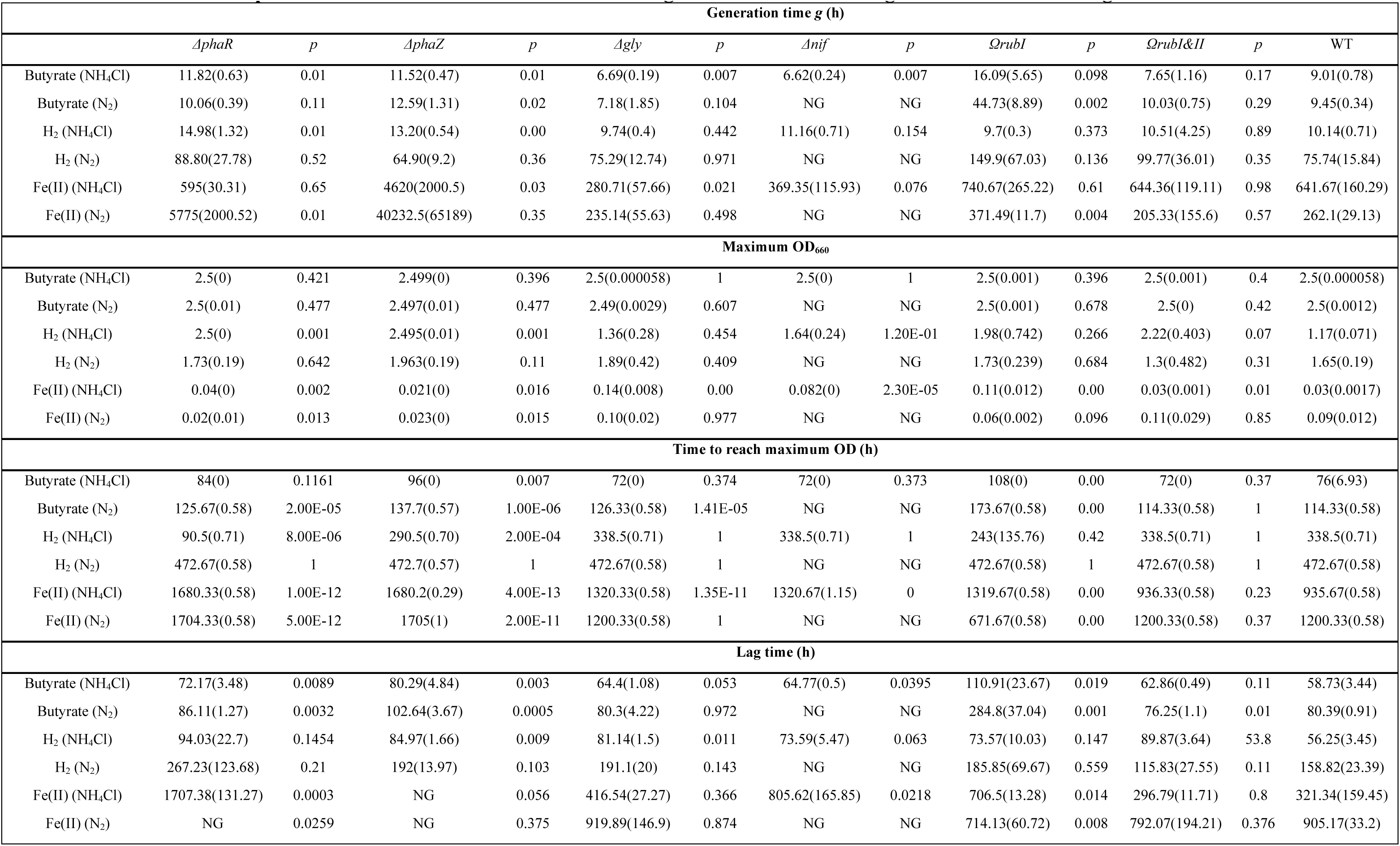
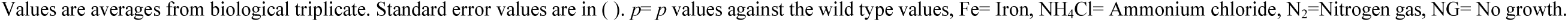
Growth parameters values of the knockouts and engineered TIE-1 strains grown under different growth conditions.

The engineered strain Ω*rub(I&II)* carrying overexpressed RuBisCO form I and II genes did not show any significant difference compared to the wild type when grown with NH_4_Cl or under nitrogen-fixing conditions (**Table 3, and** **Figure 3C** **& 3D**).

### Deletion of the *phaR* gene or overexpression of the RuBisCO form I and II increases PHB productivity in TIE-1 under photoheterotrophic growth with butyrate

We then tested the PHB productivity by all the constructed strains under photoheterotrophic growth with butyrate. Deletion of the *phaR* regulator gene increased PHB productivity approximately two-fold compared to the wild type when grown with NH_4_Cl (**Figure 3E** **& Table 4**). The remaining deletion strains (Δ*phaZ,* Δ*gly,* and Δ*nifA* double mutant) showed a decrease in PHB production compared to the wild type under both NH_4_Cl and nitrogen-fixing conditions. We observed a general trend of increased PHB production during nitrogen fixation compared to the growth with NH_4_Cl across all strains, except for Δ*phaZ*. We could not detect any PHB from the *nifA* double mutant grown under nitrogen-fixing conditions as it did not show any growth under these conditions.

**Table 4.**
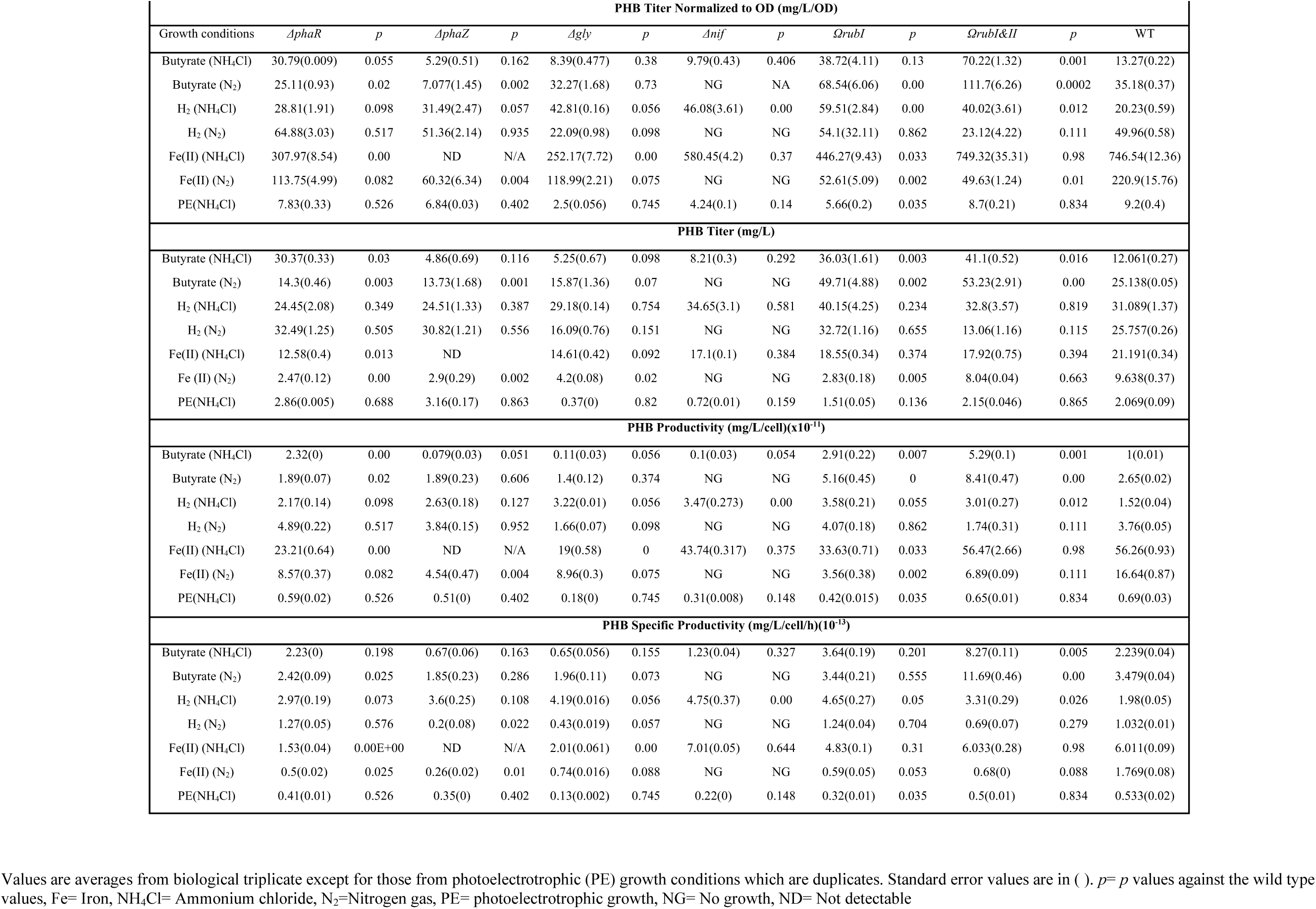
PHB production from different TIE-1 strains under photoautotrophic and heterotrophic growth conditions

The two strains harboring the two engineered Ω*rub(I)* and Ω*rub(I&II)* demonstrated increased PHB productivity. When grown photoheterotrophically with butyrate and NH_4_Cl, the Ω*rub(I)* strain had a PHB productivity twice higher than the wild type, whereas the Ω*rub(I&II*) had 5 times higher PHB productivity than the wild type. This increase of PHB was also observed under nitrogen-fixing conditions where an increase of 1.9-fold was observed from the Ω*rub(I)* and 3-fold from the Ω*rub(I&II)* (**Table 4 and** **Figure 3F**). As with the deletion mutants, an overall increase in PHB productivity was observed from the engineered and wild type strains during nitrogen fixation compared to growth with NH_4_Cl (**Figure 3F**).

### Deletion of the *phaR*, *phaZ,* or overexpression of RuBisCO form I and II increases the final optical density of TIE-1 under photoautotrophic growth conditions with H_2_ and NH_4_Cl

Achieving production from a low-cost and abundant carbon source is one of the most impactful paths in bioplastic production. Accordingly, we tested the growth of all the strains under photoautotrophic conditions using H_2_ as an electron source and CO_2_ as a carbon source in a freshwater basal liquid medium. Light is used as the energy source as TIE-1 is a photosynthetic bacterium. Growth tests were performed under nitrogen-fixing conditions with N_2_ gas or under non-nitrogen-fixing conditions with NH_4_Cl. Transitioning between heterotrophic to photoautotrophic growth conditions in H_2_ requires a metabolic shift in TIE-1 [27]. To obtain stable growth, cells were first pre-grown in yeast extract and then transferred into fresh water basal medium with H_2_ for an additional pre-growth before the growth study, and PHB extraction was performed as previously [27]. **Figure 4A** and **Table 3** show that although Δ*phaR* and Δ*phaZ* mutants showed a slightly longer generation time, they reached the maximum OD faster than the wild type when grown with H_2_ and NH_4_Cl.

**Figure 4.**
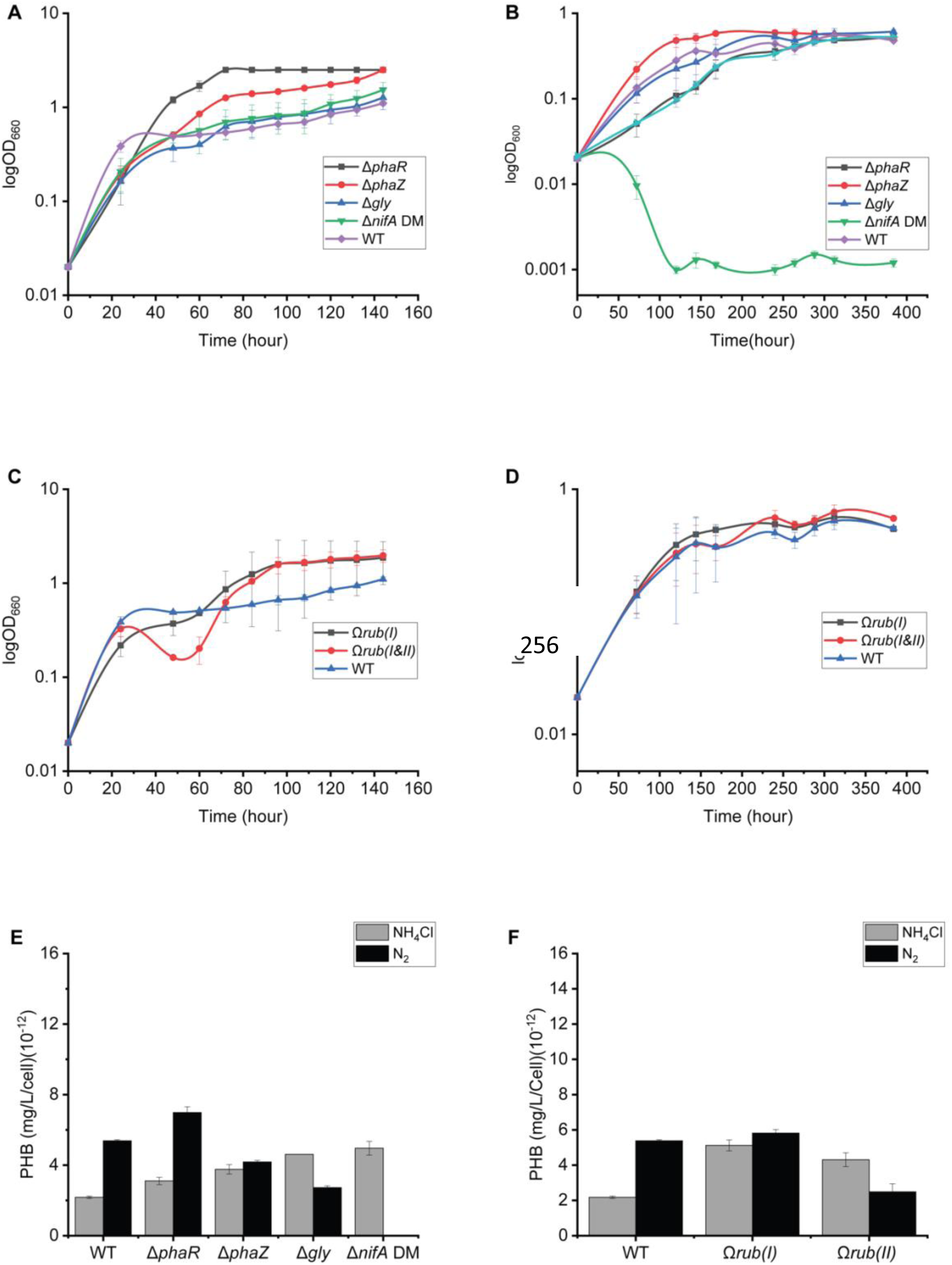
Growth and PHB productivity from different strains grown with fresh water basal media with hydrogen and NH_4_Cl or under nitrogen-fixing conditions (N_2_). **A.** Growth of all the mutant strains with NH_4_Cl. **B.** Growth of all the mutant strains under nitrogen-fixing conditions (N_2_). **C.** Growth of the engineered RuBisCO strains with NH_4_Cl. **D.** Growth of all the engineered RuBisCO stains under nitrogen-fixing conditions. **E.** PHB productivity from mutants and TIE-1 wild type strains grown under hydrogen with NH_4_Cl or N_2_. **F.** PHB productivity from Rubisco engineered and wild type strains grown under hydrogen with NH_4_Cl or N_2_. Data are from biological triplicates.

The Δ*phaR* reached the maximum OD, 3.7 times faster than the wild type whereas the Δ*phaZ* mutant reached the maximum OD almost 50 hours earlier than the wild type. The Δ*gly and* Δ*nifA* double mutant strains showed a very similar growth pattern as the wild type under growth with H_2_ and NH_4_Cl (**Figure 4A**). No significant difference was observed under nitrogen-fixing conditions from all the mutants except the Δ*nifA* double mutant where no growth was observed as shown in **Table 3** and **Figure 4B**.

Although the strains Ω*rub(I&II)* showed an extended lag time, they reached a higher final OD than the wild type (**Table 3**) when grown with H_2_ and NH_4_Cl. The Ω*rub(I)* did not show any significant growth difference compared to the wild type (**Table 3 and** **Figure 4E**). No difference was observed between the two engineered strains when growing with H_2_ under nitrogen-fixing conditions (**Table 3 and** **Figure 4D**).

### Deletion of the glycogen synthase (*gly*), *nifA* genes, or overexpression of RuBisCO form I or I&II increased PHB productivity under photoautotrophic growth with H_2_ and NH_4_Cl

We then tested the PHB productivity from all the constructed strains under growth with H_2_ under photoautotrophic conditions. The Δ*gly* and the Δ*nifA* double mutant strains showed a two-fold increase in PHB productivity when grown with H_2_ and NH_4_Cl (**Table 4 and** **Figure 4E**). The mutants Δ*phaR and* Δ*phaZ* did not show any significant difference compared to the wild type under growth with H_2_ and NH_4_Cl. Unlike growth with butyrate, switching the growth from NH_4_Cl to nitrogen increased PHB productivity in the wild type TIE-1 but not the mutant strains. In addition, all mutants showed lower PHB productivity compared to the wild type under nitrogen-fixing conditions with H_2_ as an electron source. No PHB was produced from the Δ*nifA* double mutant during the growth under nitrogen fixation conditions as no cell growth was obtained under these conditions.

Like during growth with butyrate, both engineered Ω*rub(I)* and Ω*rub(I* &*II)* strains also showed an increase in PHB productivity when grown with H_2_ and NH_4_Cl. PHB productivity obtained from both engineered strains was almost twice as what was obtained from the wild type, (**Table 4 and** **Figure 4F**) under growth with H_2_ and NH_4_Cl. While PHB productivity increased under growth with NH_4_Cl for the two engineered strains, the strain Ω*rub(I)* did not show any significant difference under nitrogen-fixing conditions when compared to the wild type (**Table 4 and** **Figure 4F**). In contrast, a decrease in PHB productivity was observed from the Ω*rub(I&II)* under the growth with H_2_ and N_2_ (**Table 4 and** **Figure 4F**).

### Deletion of the *phaR* and *phaZ* genes impaired the ability of TIE-1 to oxidize Fe(II)

One of the unique metabolisms of TIE-1 is its ability to use electrons produced by the oxidation of Fe(II) for photoautotrophy (photoferrotrophy). Under these growth conditions, the main carbon source is CO_2,_ and the energy source is light. We tested the growth of all the strains we constructed under photoferrotrophic growth conditions. All the strains were first pre-grown in H_2_ to allow expression of the genes involved in Fe(II) oxidation in TIE-1 as performed previously [27]. We observed a defect in the ability of TIE-1 to oxidize Fe(II) when the *phaZ* or *phaR* genes were deleted (**Figure 5A and 5B**.). Under growth with NH_4_Cl, the *phaZ* mutant was not able to oxidize Fe(II) even after 50 days of growth. A significant delay in Fe(II) oxidation was observed in the *phaR* mutant under growth with NH_4_Cl. The Δ*phaR* mutant was able to fully oxidize Fe(II) only after 65 days versus 40 days for the wild type. In contrast, the Δ*gly* mutants showed faster Fe(II) oxidation ability which occurred after 15 days of growth as opposed to ∼ 40 days for the wild type when grown with NH_4_Cl. The *nifA* double mutant showed similar iron oxidation pattern as the wild type under growth with NH_4_Cl (**Figure 5A**). Under nitrogen-fixing conditions, the oxidation ability of the Δ*gly* was similar to wild type TIE-1. As expected, no Fe(II) oxidation occurred from *nifA* double mutant as the strain was not able to grow under nitrogen-fixing conditions. Overexpression of RuBisCO form I appears to delay the ability of TIE-1 to oxidize Fe(II) under growth with NH_4_Cl. However, when RuBisCO form I and II are overexpressed together, the strain oxidizes Fe(II) at the same time as the wild type (∼40 days) (**Figure 5C**). During nitrogen fixation, the Ω*rubI* showed faster Fe(II) oxidation compared to the wild type, whereas the Ω*rubI&II* initiated Fe(II) oxidation at the same time as the wild type (**Figure 5D**).

**Figure 5.**
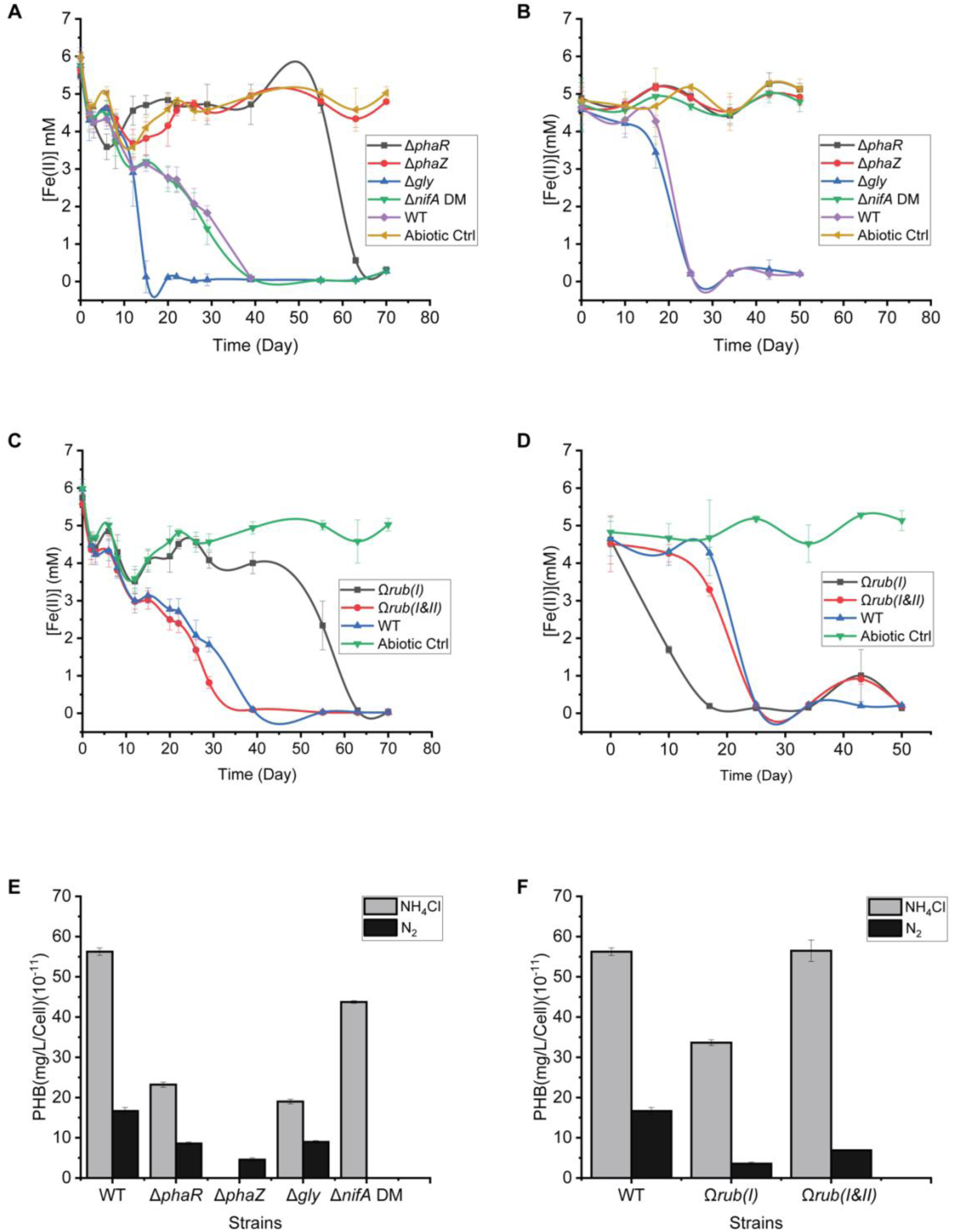
Growth and PHB productivity from strains grown under photoferrotrophy with NH_4_Cl or under nitrogen-fixing conditions N_2_. **A.** Fe(II) concentration variation from the mutants grown with NH_4_Cl. **B.** Fe(II) concentration variation from the mutans during nitrogen-fixing conditions with N_2_. **C.** Growth of the engineered RuBisCO strains with NH_4_Cl. **D.** Growth of all the engineered RuBisCO stains under nitrogen-fixing conditions **E.** PHB productivity from mutants grown with NH_4_Cl or under nitrogen fixing conditions photoferrotrophically. **F.** PHB productivity from engineered RuBisCO strains grown with NH_4_Cl or under nitrogen-fixing conditions photoferrotrophically. Data are from biological triplicates.

### Deletion of the various genes or overexpression of RuBisCO form I and II did not improve the PHB productivity in TIE-1 under photoferrotrophy

To test the PHB productivity obtained from growth during photoferrotrophy, samples were collected right after complete Fe(II) oxidation occurred. Low PHB productivity was observed across all the mutants under growth with Fe(II) and NH_4_Cl compared to the wild type (**Table 4 and** **Figure 5E**). Among all the mutants, the *nifA* double mutant had the highest PHB productivity when grown with NH_4_Cl, which was relatively close to the wild type. The Δ*phaR*, and Δ*gly* showed the lowest productivity, about half that observed in the wild type when grown with NH_4_Cl. No PHB was detected from the Δ*phaZ* mutant from the growth with NH_4_Cl which is linked to its inability to oxidize Fe(II). Like the growth with NH_4_Cl, PHB productivity for all the mutant strains during nitrogen fixation was overall, lower than that obtained from wild type (**Figure 5E and 5F**). PHB productivity obtained under nitrogen-fixing conditions was three times smaller than with NH_4_Cl for the wild type TIE-1. Δ*phaR* and Δ*gly,* mutants showed low PHB productivity compared to the wild type when grown with N_2_. PHB productivity obtained from these two mutants was almost half that of the wild type. Deleting the *phaZ* gene of TIE-1 decreased its productivity to about one-fourth of that obtained from the wild type under nitrogen-fixing conditions compared to the wild type (**Table 4 and** **Figure 5E**). No PHB was obtained from the Δ*nifA* double mutant when grown under nitrogen-fixing conditions because this strain is incapable of growth under such conditions.

Although a decrease in PHB productivity of about half was observed from the single engineered Ω*rub(I)* strain under Fe(II) and NH_4_Cl growth conditions, a similar PHB productivity to the wild type was obtained from the engineered strain carrying both RuBisCO form I and II genes (**Table 4 and** **Figure 5F**). PHB productivity obtained from the Ω*rub(I)* was about half of that obtained from the wild type under growth with Fe(II) and NH_4_Cl. Like the trend obtained from the mutants, a decrease in PHB productivity was observed in the RuBisCO overexpressing strains under nitrogen-fixing conditions compared to the growth with NH_4_Cl. PHB productivity obtained from the Ω*rub(I)* was almost 4 times less than what was obtained from the wild type under nitrogen-fixing conditions. The PHB productivity obtained from the Ω*rub(I&II)* strain dropped to 2.4 times less than the wild type under the same conditions.

### Growth under photoelectrotrophy with NH_4_Cl showed increased electron uptake from the engineered Ω*rub(I&II)* but did not reveal an increase in PHB productivity

TIE-1 has the ability to uptake electrons directly from a poised electrodes via photoelectrotrophy [29]. We were not able to calculate the growth parameters from the photoelectrotrophic of all the strains we tested because the cells did not show and increased optical density during the growth period. We then proceed to teste the ability of each of the constructed strains to uptake electrons from an electrode poised at a potential of + 100 mV vs. standard hydrogen electrode (SHE). We also measured their ability to produce PHB under growth conditions with basal media and NH_4_Cl as performed previously [27, 30, 32]. The Δ*phaZ* mutant showed a current uptake almost twice that obtained from the wild type. However, the Δ*gly* and the Δ*nifA* double mutant showed a lower current uptake compared to the wild type. A drastic decrease of around 10-fold in electron uptake was observed when the *phaR* gene was deleted from TIE-1 (**Table 5 and** **Figure 6**).

**Figure 6.**
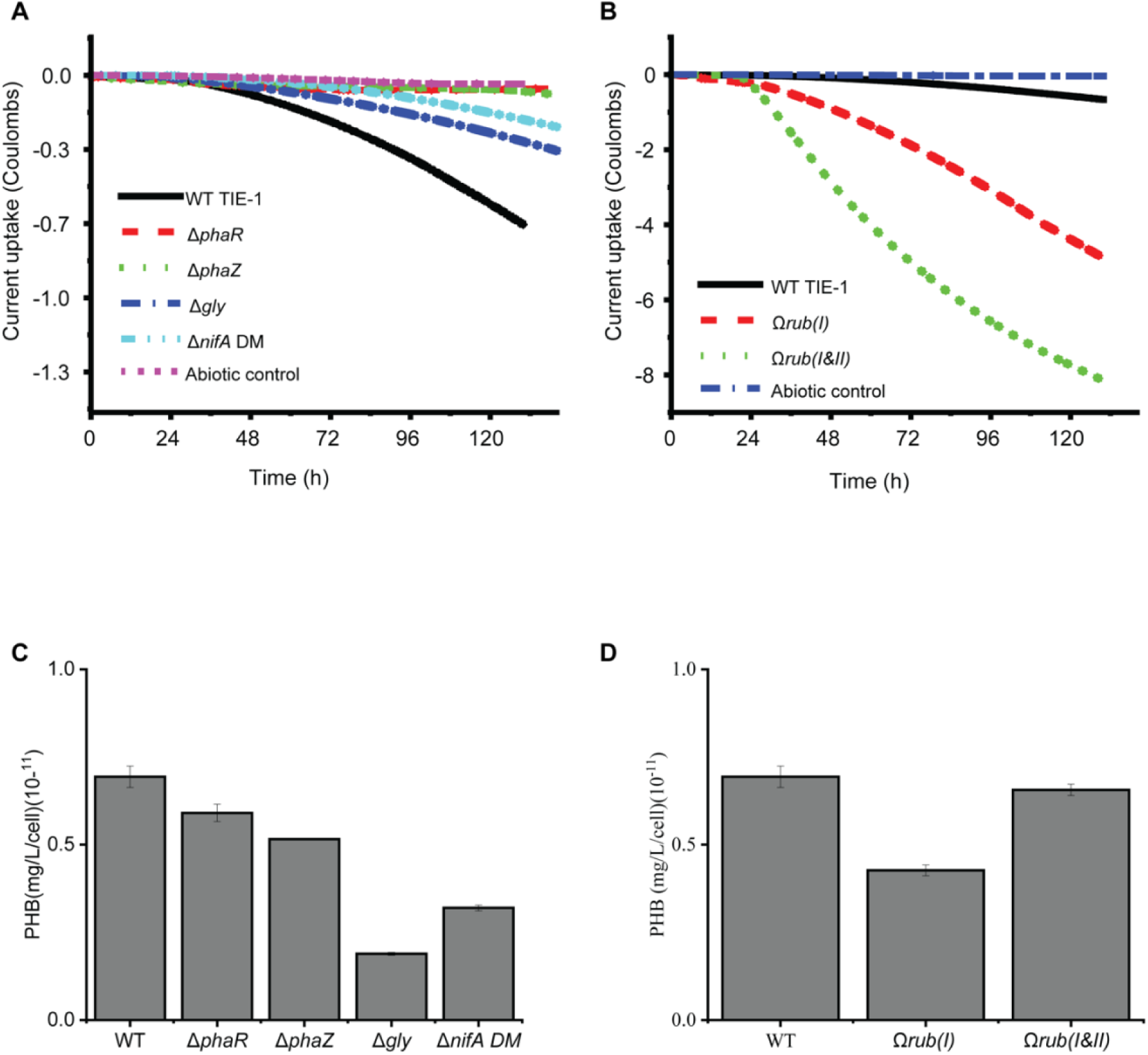
Growth and PHB production under EEU from the engineered RuBisCO strains. **A.** Current uptake from the wild type TIE-1 (WT) strain, and the various mutants Δ*phaR*, Δ*phaZ*, Δ*gly*, Δ*nifA* DM (double mutant) **B.** Current uptake from the wild type TIE-1 (WT) and the engineered strains Ω*rub(I)* and Ω*rub(I&II)*. **C.** PHB productivity from mutants grown with NH_4_Cl photoelectrotrophically. **D.** PHB productivity from engineered RuBisCO strains grown with NH_4_Cl under photoelectrotrophic growth conditions. Data are from biological duplicate.

**Table 5.** Current uptake and current density obtained from the different strains during photoelectroautotrophic growth with NH_4_Cl.

| Stains | Current density ($\mu A/cm^2$) | Current uptake (C) | *p* |
| --- | --- | --- | --- |
| $\Delta phaR$ | $-0.0729 \pm 0.0019$ | -0.0559 | 0.00 |
| $\Delta phaZ$ | $-0.6735 \pm 0.0326$ | -1.2284 | 0.00 |
| $\Delta gly$ | $-1.1131 \pm 0.034$ | -0.4588 | 0.00 |
| $\Delta nifA$ DM | $-0.9029 \pm 0.0777$ | -0.3288 | 0.00 |
| $\Omega rub(I)$ | $-17.6725 \pm 0.7832$ | -4.8896 | 0.00 |
| $\Omega rub(I&II)$ | $-31.5573 \pm 0.8221$ | -8.1316 | 0.00 |
| Blank | $0.0059 \pm 0.0103$ | -0.0384 | 0.00 |
| WT TIE-1 | $-2.66759 \pm 0.1093$ | -0.6648 | 0.00 |

Values are from biological replicates, *p* - *p* values vs. wild type.

An increase in current uptake was observed in the two engineered RuBisCO strains compared to the wild type. An increase of almost ten and eight-fold was obtained from the Ω*rub(I&II)* and Ω*rub(I)* respectively (**Table 5**, **Figure 6**) compared to the wild type. We then tested the PHB productivity of the different strains under the photoelectrotrophic growth conditions. The lower electron uptake from the mutants Δ*phaR,* Δ*gly,* and the Δ*nifA* double mutant was accompanied by a lower PHB productivity compared to the wild type. The PHB productivity obtained from the Δ*gly* was the lowest amongst all the mutants, with around three times lower PHB production than the wild type. This was followed by the *nifA* double mutant which was almost half of the wild type. The PHB productivity obtained from Δ*phaR* was similar to the wild type. Overall, none of the mutants showed a PHB productivity higher than the wild type (**Table 5 and** **Figure 6C**). Similarly, no noticeable increase in PHB productivity was obtained from the engineered RuBisCO strains. The PHB obtained from the Ω*rub(I&II)* was very similar to the productivity obtained from the wild type whereas the Ω*rub(I)* was 25% lower than the wild type (**Table 5**).

## Discussion

In the present study we show that TIE-1 can be modified using different genetic engineering and synthetic biology tools to improve PHB production. Both deletion of competitive pathways and regulatory genes of the PHB cycle increase PHB productivity. Also, TIE-1 overexpressing RuBisCO form I and II showed increased PHB productivity. This shows that our newly developed φC31 integration system is effective and useful for genetic manipulation in TIE-1.

### Integration system in TIE-1

We explored two φC31 integration systems: inducible control of φC31 integrase, and constitutive expression of φC31 integrase. The inducible system used a self-replicating plasmid while the constitutive system used a suicide plasmid. This design was intended to ensure that optimal expression of φC31 integrase is achieved. As a consequence, the inducible system will leave a plasmid behind, while the constitutive system will not. We observed that the genome-based system with inducible control of φC31 integrase is the most efficient; however, integrating φC31 into the genome relies on two-step integration. Thus, this process is harder to apply to other organisms. On the other hand, plasmid-based systems are much easier to apply to other closely related systems. Between the two plasmid-based systems, using a constitutive promoter to drive φC31integrase expression resulted in higher efficiency and did not leave a plasmid behind because it uses a suicide plasmid. Thus, the plasmid-based system with the constitutive promoter is easy to use and demonstrates relatively high efficiency.

### Effect of the deletion of genes from TIE-1 on its growth and PHB productivity

Deletion of the *phaR* resulted in a growth defect in TIE-1 both under photoheterotrophic with butyrate and photoautotrophic with H_2_. It also decreased its ability to oxidize Fe(II), and uptake electrons directly from the electrode, decreased its PHB productivity compared to the wild type. Inactivation of the *phaR* in *Bradyrhizobium diazoefficiens* USDA110 not only decreased PHB accumulation but also affected other processes such as exopolysaccharide (EPS) biosynthesis, and heat stress tolerance [63]. This suggests that the role of PhaR in *Bradyrhizobium diazoefficiens* USDA110 is not limited to PHB biosynthesis only. In TIE-1, comparing Δ*phaR* to WT using all the growth and PHB parameters as shown in **Tables 3 and 4**, we are unable to conclude what the loss of PhaR in in TIE-1 and how it affects PHB production. Comparing expression changes in the PHB cycle genes and also, differences in the PHB enzyme abundance in the Δ*phaR* vs. WT will be informative.

The purpose of deleting the *phaZ* gene of TIE-1 was to prevent depolymerization of PHB polymer, which should result in its accumulation. We observed that the deletion of *phaZ* in TIE-1 overall did not significantly increase PHB productivity in TIE-1. However, it is worth mentioning that deletion of *phaZ* affected growth of TIE-1 under photoheterotrophic with butyrate both with NH_4_Cl and under nitrogen-fixing conditions. We also observed a defect in the ability of the Δ*phaZ* mutant to oxidize Fe(II) during photoferrotrophic growth both with NH_4_Cl and under nitrogen-fixing conditions. Because PHB is considered an electron sink, deletion of the PhaZ might have caused an electron imbalance in TIE-1 [38].

Deletion of the glycogen synthase (*gly*) gene did not affect the growth of TIE-1 under photoheterotrophic growth with butyrate or under photoautotrophic growth with H_2_, Fe(II) or with a poised electrode. However, a decrease in PHB productivity was observed under all photoheterotrophic growth with butyrate and photoautotrophic growth conditions under nitrogen-fixing conditions. This effect of the deletion of the glycogen synthase gene on PHB accumulation is similar to what was discovered previously in *Synechocystis sp*. PCC 6803. Here glycogen was hypothesized to be a carbon source for PHB synthesis under nitrogen (NaNO_3_) starvation [50]. This could explain the increase in PHB productivity under photoautotrophic growth conditions with H_2_ and NH_4_Cl.

### Effect of overexpressing RuBisCO genes on TIE-1’s growth and PHB productivity

We observed an increase of PHB productivity in TIE-1 overexpressing RuBisCO form I and form I&II during photoheterotrophic growth with butyrate and photoautotrophically with H_2_ (**Table 4 and** **Figure 3F** **& 4F**). An increase in carbon fixation in the CBB cycle is expected to increase the abundance of acetyl-CoA [64], the precursor of PHB. We observed an increase in PHB productivity, which could be a direct result of overexpression of the two RuBisCO genes. Overexpression of the RuBisCO gene in *Ralstonia eutropha* (now *C. necator*) grown autotrophically with minimum media resulted in a 99.7% increase in PHB accumulation [54]. We achieved an increase of an overall 2-fold PHB productivity from the Ω*rub(I)* both under photoheterotrophic with butyrate (both with NH_4_Cl and N_2_) and photoautotrophic with H_2_ and NH_4_Cl compared to wild type. The Ω*rub(I&II)* strains reached an even higher PHB productivity of 5-fold and 3-fold under photoheterotrophic growth with butyrate and NH_4_Cl, and with butyrate under nitrogen-fixing conditions compared to wild type, respectively. We observed a cumulative effect of overexpression both RuBisCO form I and form II in increasing PHB productivity under photoheterotrophic growth with butyrate. An increase of 2-fold in PHB productivity was also observed in the Ω*rub(I&II)* under photoautotrophic growth conditions with H_2_ and NH_4_Cl. Surprisingly, the Ω*rub(I)* showed a growth defect when grown photoheterotrophically with butyrate. Moreover, the Ω*rub(I)* had a defect in its ability to oxidize Fe(II) when grown photoferrotrophically with Fe(II) and NH_4_Cl. This defect in Fe(II) oxidation might have caused the decrease in PHB productivity from the Ω*rub(I)* under photoferrotrophy. The defect in PHB productivity in Ω*rub(I)* seems to disappear when both RuBisCO forms I and II (Ω*rub(I&II)* are overexpressed in TIE-1 regardless of the nitrogen source.

### Effect of deletion of the nifA genes on PHB productivity and growth of TIE-1

Nitrogen fixation has been previously reported to increase PHB production [27, 38, 52]. Our results confirm that switching TIE-1 to fix dinitrogen gas increases PHB productivity under photoheterotrophic growth conditions with butyrate. Under photoautotrophy with H_2_ this trend was not fully conserved. This also agrees with our previous findings where the increase in PHB productivity observed under heterotrophic growth was lost when cells were switched to autotrophic growth conditions [27]. Deletion of the *nifA* genes decreased PHB productivity under photoheterotrophic growth with butyrate and NH_4_Cl or with N_2_. Similar to the increase in *n*-butanol production observed in the Δ*nifA* double mutant TIE-1 strain carrying the five-butanol gene cassette[34], we noticed an increase in PHB productivity from Δ*nifA* double mutant strain grown under photoautotrophy with H_2_ and NH_4_Cl [34]. This increase in PHB productivity from the Δ*nifA* double mutant strain was only limited to this growth condition even though we expected to see an increase of PHB productivity from across all growth conditions we tested. We observed that while all the mutant strains showed a decrease in PHB production under photoautotrophic growth with Fe(II) and NH_4_Cl, the Δ*nifA* double mutant’s PHB productivity remained relatively close to the wild type (**Table 4 and** **Figure 5E**).

### Future directions

We conclude that overexpressing the RuBisCO form I and II using the φC31 system increased PHB productivity in TIE-1. We also observed that deletion of *phaR* under photoheterotrophic growth with butyrate and NH_4_Cl increased the PHB productivity. In addition, deletion of the glycogen synthase (*gly*) or *nifA* regulators increases PHB productivity under photoautotrophic growth with H_2_ and NH_4_Cl. It would be interesting to assess the effect of combining the deletion of *phaR*, *phaZ*, *gly,* and the *nifA* regulators genes with the overexpression of the RuBisCO form I and II genes in TIE-1.

## Materials and Methods

### Bacterial strains, media, and growth conditions

**Table 1** lists all the strains used in the study. Lysogeny broth (LB) was used for the growth of all *E. coli* strains at 37°C. *Rhodopseudomonas palustris* TIE-1was grown in a medium containing 3 g/L yeast extract, 3 g/L peptone, 10 mM MOPS [3-N (morpholino) propanesulphonic acid] (pH 7.0), and 10 mM succinate (YPSMOPS) at 30 °C. LB and YPSMOPS agar plates were prepared with the addition of 15g/L agar. When needed, the antibiotic or sucrose was added as indicated in **Table S1**. All *E. coli* strains were grown on Lysogeny Broth (LB) at 37 degrees C. TIE-1 was grown in YPSMOPS, a medium composed of 3 g/L yeast extract, 3 g/L peptone, 10 mM 3-N morpholino propane sulphonic acid (MOPS), and 10 mM succinate at 30 degrees C. Plates were made with LB and 15 g/L agar or YPSMOPS and 15 g/L agar. Table 1 contains a list of the strains used in the study. **Table S1** shows the concentration of antibiotics used as positive and negative selection components.

### Plasmid construction

All the plasmids used in this study are listed in **Table 2**. The kanamycin and chloramphenicol gene sequences were PCR amplified from pSRKKm and pSRKCm, respectively. All these antibiotic resistance marker genes were then cloned into pJQ200KS plasmid separately to replace the gentamicin resistance gene resulting in pWB091 and pWB092. All the primers used are listed in **Table S2.**

### Mutant construction

The φC31 *attB* and *attP* sequences are obtained from the previously published sequences [59]. The *attP* sequence was cloned into pJQ200KS, resulting in pWB081. Then the *P_aphII_*-*mCherry*-fd cassette was cloned into pJQ200KS, resulting in pWB081. For the plasmid-based system, the φC31 integrase sequence was cloned to either plasmid pSRKGm or pWB081, resulting in pWB084 and pWB088. For the genome-based system, the φC31 integrase sequence was cloned to pAB314, resulting in pWB089. The *attB* sequence was also cloned into pAB314, resulting in pWB083.

### Construction of engineered TIE-1

Strains were constructed as described in Bose et al. (2011)[31]. pWB107 and pWB108 were individually conjugated into TIE-1 through *E. coli* S17-1/λ. After two successive homologous recombination, successful integrants were screened by PCR as shown in **Figure S4**.

### TIE-1 electroporation

To prepare electrocompetent cells, the TIE-1 strain was inoculated in 500 mL YPSMOPS and then incubated at 30 degree C. After reaching an OD_660_ of 0.5∼0.6, the culture was centrifuged and washed at 4 degree C at 4000 X *g* for 10 minutes. After five washes with 10% glycerol, the cell pellet was resuspended in only 2 mL of ice-cold 10% glycerol. This resuspension was aliquoted by 50 μL per sample then saved in -80 degree C. For every electroporation, 0.2 μg of plasmid was added to 50 μL thawed electrocompetent cells and mixed well. This mixture was added to 1mm gap cuvettes and then electroporated at 1.8kV using a Biorad gene Micropulser. After electroporation the pulsed mixture was added to 2 mL warmed Super Optimal Broth (SOB) or LB and grown for 45 minutes. 10 μL, 100μL, 500μL, and the remainder of the culture were plated on a selective medium. Because these plates will be further imaged by Nikon A1, glass Petri dishes instead of plastic Petri dishes were used.

### Imaging of *mCherry*

After 4 to 5 days of incubation, the plated electroporated culture was imaged by Nikon A1 confocal Microscope. For each plate, an image of the whole plate was captured using a camera through both the bright field and the Texas-red channel with a 10X scope. The colony number of each plate was quantified using NIS-Elements AR Analysis 5.11.01 64-bit software.

### Calculations of transformation efficiency and Colony Forming Unit per Optical Density

We evaluated our phage integration systems by calculating the transformation efficiency as calculated by the following equation:

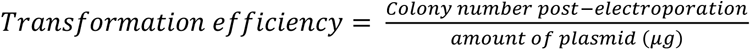

The editing efficiency is calculated by:

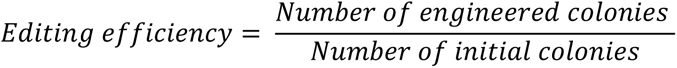

### Construction of TIE-1 mutants

The following strains were constructed using the markerless deletion method as described in **Figure S1**: *phaZ* mutant ΔRpal_0578, and *phaR* mutant (ΔRpal_0531). The two mutants Δ*gly* and the Δ*nifA* double mutant have been described previously [34]. The mutant constructions were as follows: 1kb upstream and 1kb downstream of each of the genes of interest were PCR amplified and cloned into the cloning vector pJQ200KS. The constructed plasmids were then electroporated into the donor strain *E. coli* S17-1/λ. The *E. coli* donor strains carrying the constructed plasmids were conjugated with the TIE-1 wild type. After two consecutive homologous recombination events, successful mutant candidates were screened using PCR (**Figure S2**) and the primers listed in **Table S3**.

### PHB extraction and analysis

PHB samples were extracted and analyzed as previously reported [27]. Briefly, 5 mL of liquid bacterial samples. PHB from the pelleted samples was extracted using methanol and chloroform of 325 µL and 400 µL respectively. Simultaneously, PHB polymers were digested into its monomer crotonic acid using 75 µL. The mixtures were incubated at 95°C for one hour. Phase separation and acid elimination were done by the addition of 500 µL of LC-MS grade water. The organic phases were dried using a speed vacuum. Dried crotonic acid was then resuspended in 50% acetonitrile/50% water and analyzed using an Agilent Technologies 6420 Triple Quad LC/MS. Crotonic acids were detected with mass to charge ratio (*m/z*)=87.

## Acknowledgments

We would like to thank Dr. Joshua Blodgett for facilitating the PHB analysis using LC-MS. We greatly appreciate the contribution of Eric Conners in proofreading the manuscript. This work was supported by the following grants to A.B.: The David and Lucile Packard Foundation Fellowship (201563111), the U.S. Department of Energy (grant number DESC0014613), and the U.S. Department of Defense, Army Research Office (grant number W911NF-18-1-0037), Gordon and Betty Moore Foundation, National Science Foundation (Grant Number 2021822, Grant Number 2124088, and Grant Number 2117198), the U.S. Department of Energy by Lawrence Livermore National Laboratory under Contract DEAC5207NA27344 (LLNL-JRNL-812309), an NIGMS grant (NIHR01GM141344), and a DEPSCoR grant (FA9550-21-1-0211). A.B. was also funded by a Collaboration Initiation Grant, an Office of the Vice-Chancellor of Research Grant, an International Center for Energy, Environment, and Sustainability Grant and a SPEED grant from Washington University in St. Louis.

## Author contributions

TOR, WB, MS, JO, and HS performed all the experiments. RK assisted with the bioelectrochemical experiments. TOR, WB, and AB wrote the manuscript with input from all authors.

## References

1. Keasling, J.D., Manufacturing Molecules Through Metabolic Engineering. Science, 2010. 330(6009): p. 1355.

2. Müller, T., et al., Synthetic mutualism in engineered E. coli mutant strains as functional basis for microbial production consortia. Engineering in Life Sciences, 2023. 23(1): p. e2100158.

3. Panda, S. and K. Zhou, Engineering microbes to overproduce natural products as agrochemicals. Synthetic and Systems Biotechnology, 2023. 8(1): p. 79–85.

4. Kyriakou, M., et al., Enhancing bioproduction and thermotolerance in *Saccharomyces cerevisiae* via cell immobilization on biochar: Application in a citrus peel waste biorefinery. Renewable Energy, 2020.

5. Saini, M., et al., Potential production platform of n-butanol in *Escherichia coli*. Metabolic engineering, 2015. 27: p. 76–82.

6. Wang, J., et al., A novel process for cadaverine bio-production using a consortium of two engineered *Escherichia coli*. Frontiers in microbiology, 2018. 9: p. 1312.

7. Li, H., et al., Effects of metabolic pathway gene copy numbers on the biosynthesis of (2S)- naringenin in *Saccharomyces cerevisiae*. Journal of Biotechnology, 2020.

8. Orozco, P.A., Lanthipeptide biosynthesis in marine *Synechococcus*: expression, characterization, and bioengineering of a wide variety of synechococsins. 2023.

9. Zhang, F., S. Rodriguez, and J.D. Keasling, Metabolic engineering of microbial pathways for advanced biofuels production. Current Opinion in Biotechnology, 2011. 22(6): p. 775–783.

10. Miller, E.M. and J.A. Nickoloff, *Escherichia coli* electrotransformation, in Electroporation protocols for microorganisms. 1995, Springer. p. 105–113.

11. Mäkelä, J. and D.J. Sherratt, Organization of the *Escherichia coli* chromosome by a MukBEF axial core. Molecular cell, 2020.

12. Van Rossum, H.M., et al., Engineering cytosolic acetyl-coenzyme A supply in *Saccharomyces cerevisiae*: pathway stoichiometry, free-energy conservation and redox-cofactor balancing. Metabolic engineering, 2016. 36: p. 99–115.

13. DiCarlo, J.E., et al., Genome engineering in *Saccharomyces cerevisiae* using CRISPR-Cas systems. Nucleic acids research, 2013. 41(7): p. 4336–4343.

14. Çakar, Z.P., et al., Evolutionary engineering of *Saccharomyces cerevisiae* for improved industrially important properties. FEMS yeast research, 2012. 12(2): p. 171–182.

15. Den Haan, R., et al., Engineering *Saccharomyces cerevisiae* for next generation ethanol production. Journal of Chemical Technology & Biotechnology, 2013. 88(6): p. 983–991.

16. Abbott, D.A., et al., Metabolic engineering of *Saccharomyces cerevisiae* for production of carboxylic acids: current status and challenges. FEMS yeast research, 2009. 9(8): p. 1123–1136.

17. Cao, X., et al., Engineering yeast for high-level production of diterpenoid sclareol. Metabolic Engineering, 2023. 75: p. 19–28.

18. Parapouli, M., et al., *Saccharomyces cerevisiae* and its industrial applications. AIMS microbiology, 2020. 6(1): p. 1.

19. Luo, H., et al., Efficient bio-butanol production from lignocellulosic waste by elucidating the mechanisms of *Clostridium acetobutylicum* response to phenolic inhibitors. Science of The Total Environment, 2020. 710: p. 136399.

20. Wang, Q., et al., Complete PHB mobilization in *Escherichia coli* enhances the stress tolerance: a potential biotechnological application. Microbial cell factories, 2009. 8(1): p. 1–9.

21. Ferreira, S., et al., Metabolic engineering strategies for butanol production in *Escherichia coli*. Biotechnology and Bioengineering, 2020. 117(8): p. 2571–2587.

22. Gong, C., et al., Genetic manipulation strategies for ethanol production from bioconversion of lignocellulose waste. Bioresource Technology, 2022: p. 127105.

23. Calero, P. and P.I. Nikel, Chasing bacterial chassis for metabolic engineering: a perspective review from classical to non-traditional microorganisms. Microbial Biotechnology, 2019. 12(1): p. 98–124.

24. Azambuja, S.P. and R. Goldbeck, Butanol production by *Saccharomyces cerevisiae*: perspectives, strategies and challenges. World Journal of Microbiology and Biotechnology, 2020. 36(3): p. 1–9.

25. Brown, B., M. Wilkins, and R. Saha, *Rhodopseudomonas palustris*: A biotechnology chassis. Biotechnology Advances, 2022: p. 108001.

26. Jiao, Y., et al., Isolation and Characterization of a Genetically Tractable Photoautotrophic Fe(II)- Oxidizing Bacterium, *Rhodopseudomonas palustris* Strain TIE-1. Applied and Environmental Microbiology, 2005. 71(8): p. 4487–4496.

27. Ranaivoarisoa, T.O., et al., Towards sustainable bioplastic production using the photoautotrophic bacterium *Rhodopseudomonas palustris* TIE-1. Journal of industrial microbiology & biotechnology, 2019. 46(9-10): p. 1401–1417.

28. Karthikeyan, R., R. Singh, and A. Bose, Microbial electron uptake in microbial electrosynthesis: a mini-review. Journal of industrial microbiology & biotechnology, 2019. 46(9-10): p. 1419–1426.

29. Bose, A., et al., Electron uptake by iron-oxidizing phototrophic bacteria. Nat Commun 5: 3391. 2014.

30. Rengasamy, K., et al., An insoluble iron complex coated cathode enhances direct electron uptake by *Rhodopseudomonas palustris TIE-1*. Bioelectrochemistry, 2018. 122: p. 164–173.

31. Bose, A. and D.K. Newman, Regulation of the phototrophic iron oxidation (pio) genes in *Rhodopseudomonas palustris* TIE-1 is mediated by the global regulator, FixK. Molecular Microbiology, 2011. 79(1): p. 63–75.

32. Rengasamy, K., et al., Magnetite nanoparticle anchored graphene cathode enhances microbial electrosynthesis of polyhydroxybutyrate by *Rhodopseudomonas palustris* TIE-1. Nanotechnology, 2020. 32(3): p. 035103.

33. Rabaey, K., P. Girguis, and L.K. Nielsen, Metabolic and practical considerations on microbial electrosynthesis. Current opinion in biotechnology, 2011. 22(3): p. 371–377.

34. Bai, W., et al., n-Butanol production by *Rhodopseudomonas palustris* TIE-1. Communications Biology, 2021. 4(1): p. 1257.

35. Gupta, D., et al., Photoferrotrophs Produce a PioAB Electron Conduit for Extracellular Electron Uptake. mBio, 2019. 10.

36. Singh, R., et al., Genetic redundancy in iron and manganese transport in the metabolically versatile bacterium *Rhodopseudomonas palustris* TIE-1. Applied and Environmental Microbiology, 2020.

37. Guzman, M.S., et al., Phototrophic extracellular electron uptake is linked to carbon dioxide fixation in the bacterium *Rhodopseudomonas palustris*. Nature communications, 2019. 10(1): p. 1–13.

38. McKinlay, J.B., et al., Non-growing *Rhodopseudomonas palustris* increases the hydrogen gas yield from acetate by shifting from the glyoxylate shunt to the tricarboxylic acid cycle. Journal of Biological Chemistry, 2014. 289(4): p. 1960–1970.

39. Gosse, J.L., et al., Hydrogen production by photoreactive nanoporous latex coatings of nongrowing *Rhodopseudomonas palustris* CGA009. Biotechnology progress, 2007. 23(1): p. 124–130.

40. Meereboer, K.W., M. Misra, and A.K. Mohanty, Review of recent advances in the biodegradability of polyhydroxyalkanoate (PHA) bioplastics and their composites. Green Chemistry, 2020. 22(17): p. 5519–5558.

41. Sabbagh, F. and I.I. Muhamad, Production of poly-hydroxyalkanoate as secondary metabolite with main focus on sustainable energy. Renewable and Sustainable Energy Reviews, 2017. 72: p. 95–104.

42. Karan, H., et al., Green Bioplastics as Part of a Circular Bioeconomy. Trends in Plant Science, 2019. 24(3): p. 237–249.

43. Bátori, V., et al., Anaerobic degradation of bioplastics: A review. Waste Management, 2018. 80: p. 406–413.

44. Sirohi, R., et al., Critical overview of biomass feedstocks as sustainable substrates for the production of polyhydroxybutyrate (PHB). Bioresource Technology, 2020. 311: p. 123536.

45. Brodin, M., et al., Lignocellulosics as sustainable resources for production of bioplastics–A review. Journal of Cleaner Production, 2017. 162: p. 646–664.

46. Kourmentza, C., et al., Recent advances and challenges towards sustainable polyhydroxyalkanoate (PHA) production. Bioengineering, 2017. 4(2): p. 55.

47. Maehara, A., et al., PhaR, a protein of unknown function conserved among short-chain-length polyhydroxyalkanoic acids producing bacteria, is a DNA-binding protein and represses *Paracoccus denitrificans* phaP expression in vitro. FEMS microbiology letters, 2001. 200(1): p. 9–15.

48. York, G.M., J. Stubbe, and A.J. Sinskey, The *Ralstonia eutropha* PhaR protein couples synthesis of the PhaP phasin to the presence of polyhydroxybutyrate in cells and promotes polyhydroxybutyrate production. Journal of bacteriology, 2002. 184(1): p. 59–66.

49. Cervenka, E., Optimizing PHB Bioplastic Accumulation in *Sinorhizobium meliloti* grown on CO_2_-derived Formate and Bicarbonate. 2022.

50. Koch, M., et al., PHB is produced from glycogen turn-over during nitrogen starvation in *Synechocystis sp. PCC 6803*. International journal of molecular sciences, 2019. 20(8): p. 1942.

51. Wang, J. and H.-Q. Yu, Biosynthesis of polyhydroxybutyrate (PHB) and extracellular polymeric substances (EPS) by *Ralstonia eutropha* ATCC 17699 in batch cultures. Applied microbiology and biotechnology, 2007. 75: p. 871–878.

52. Higuchi-Takeuchi, M., et al., Synthesis of high-molecular-weight polyhydroxyalkanoates by marine photosynthetic purple bacteria. Plos one, 2016. 11(8): p. e0160981.

53. Kamravamanesh, D., et al., Photosynthetic poly-β-hydroxybutyrate accumulation in unicellular cyanobacterium *Synechocystis sp. PCC 6714*. AMB express, 2017. 7(1): p. 1–12.

54. Li, Z., et al., Engineering the Calvin–Benson–Bassham cycle and hydrogen utilization pathway of *Ralstonia eutropha* for improved autotrophic growth and polyhydroxybutyrate production. Microbial cell factories, 2020. 19(1): p. 1–9.

55. Guss, A.M., et al., New methods for tightly regulated gene expression and highly efficient chromosomal integration of cloned genes for *Methanosarcina* species. Archaea (Vancouver, B.C.), 2008. 2(3): p. 193–203.

56. Huff, J., et al., Taking phage integration to the next level as a genetic tool for mycobacteria. Gene, 2010. 468(1): p. 8–19.

57. Huang, H., et al., Phage serine integrase-mediated genome engineering for efficient expression of chemical biosynthetic pathway in gas-fermenting *Clostridium ljungdahlii*. Metabolic Engineering, 2019. 52: p. 293–302.

58. Baltz, R.H., Streptomyces temperate bacteriophage integration systems for stable genetic engineering of actinomycetes (and other organisms). Journal of industrial microbiology & biotechnology, 2012. 39(5): p. 661–672.

59. Groth, A.C., et al., A phage integrase directs efficient site-specific integration in human cells. Proceedings of the national academy of Sciences, 2000. 97(11): p. 5995–6000.

60. Quandt, J. and M.F. Hynes, Versatile suicide vectors which allow direct selection for gene replacement in Gram-negative bacteria. Gene, 1993. 127(1): p. 15–21.

61. Khan, S.R., et al., Broad-host-range expression vectors with tightly regulated promoters and their use to examine the influence of TraR and TraM expression on Ti plasmid quorum sensing. Applied and environmental microbiology, 2008. 74(16): p. 5053–5062.

62. Phillips, G.J., New cloning vectors with temperature-sensitive replication. Plasmid, 1999. 41(1): p. 78–81.

63. Nishihata, S., et al., *Bradyrhizobium diazoefficiens* USDA110 PhaR functions for pleiotropic regulation of cellular processes besides PHB accumulation. BMC microbiology, 2018. 18(1): p. 1–17.

64. Miao, R., et al., Current processes and future challenges of photoautotrophic production of acetyl-CoA-derived solar fuels and chemicals in cyanobacteria. Current opinion in chemical biology, 2020. 59: p. 69–76.

