## Supplemental Materials for "Improving bioplastic production by *Rhodopseudomonas palustris* TIE-1 using synthetic biology and metabolic engineering"

### Supplementary Figures and Tables:

**Figure S1.**

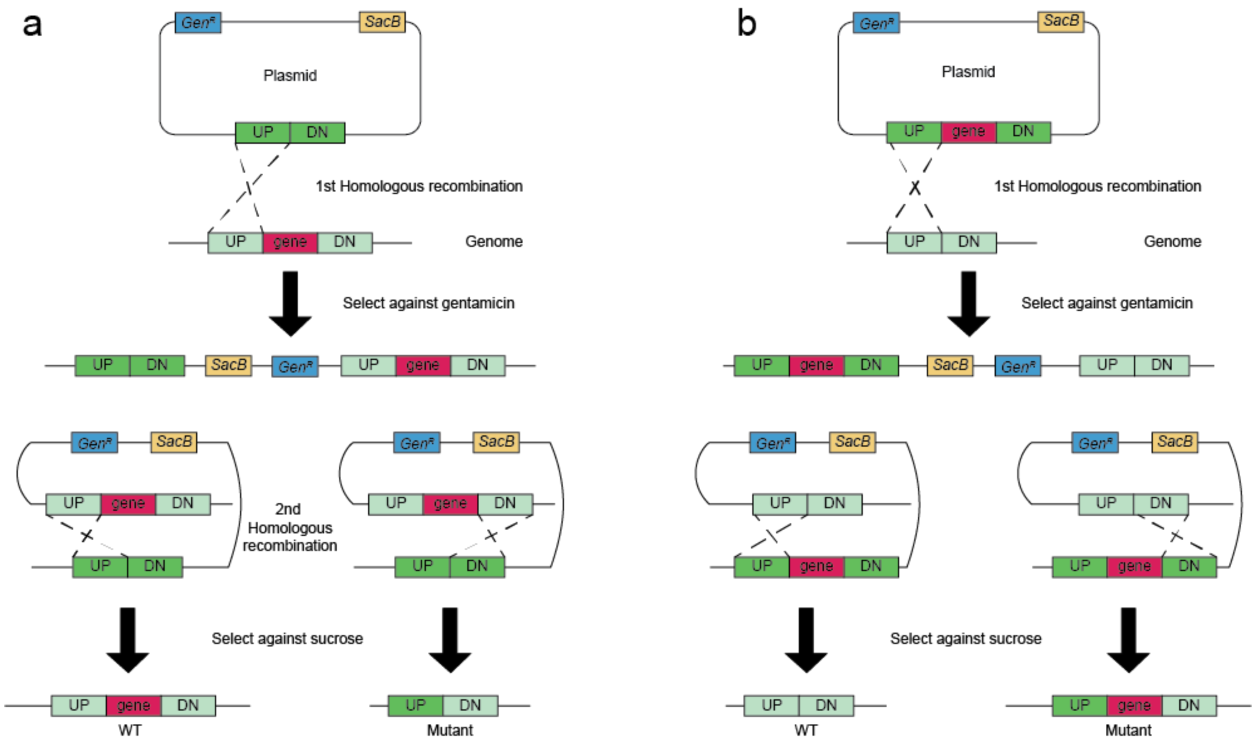


**Figure S1.** Schematic flow of two-step integration. a. Schematic flow of generating a knockout mutant b. Schematic flow of generating a knock-in mutant UP: upstream homologous arm, DN: down-stream homologous arm, Gen$^r$: gentamicin resistance, *SacB*: sucrose counter-selection marker.

**Figure S2.**

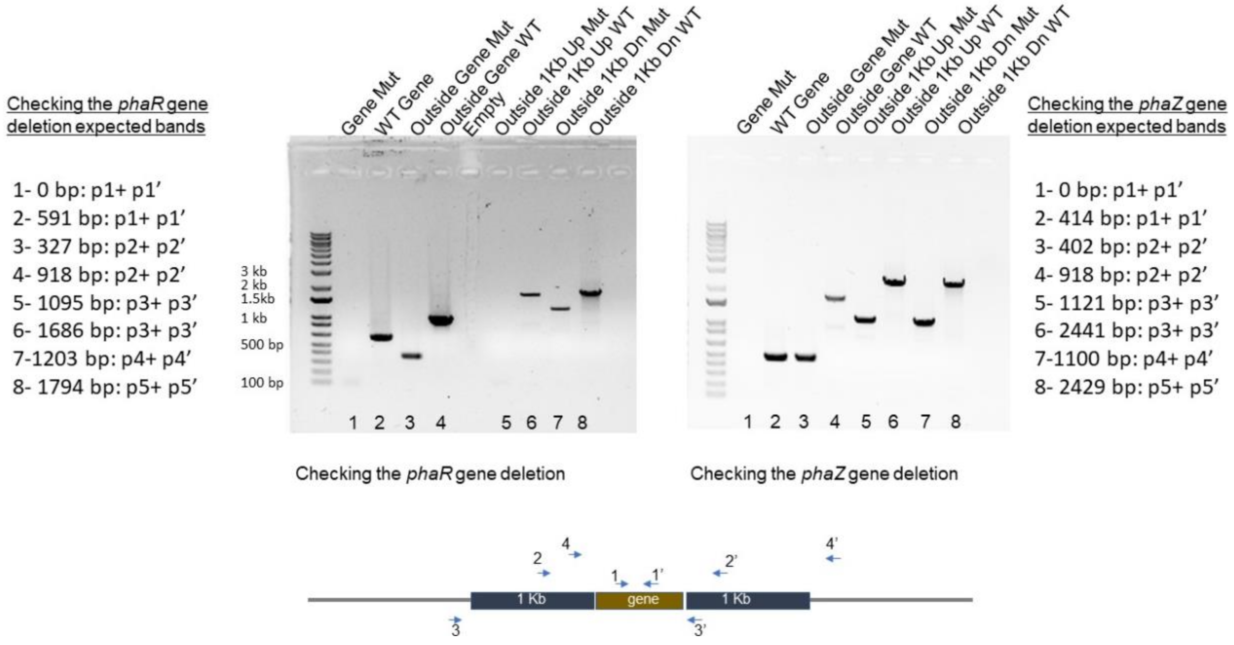


**Figure S2.** Checking the Δ*phaR* and Δ*phaZ* mutants using PCR amplification. The primers set used for the PCR test are shown at the bottom of the figures. Left side of the gel images show the expected sizes from the primers sets used for checking the Δ*phaR* mutant. Right side of the gel images show the expected sizes bands from the Δ*phaZ* mutant.

**Figure S3.**

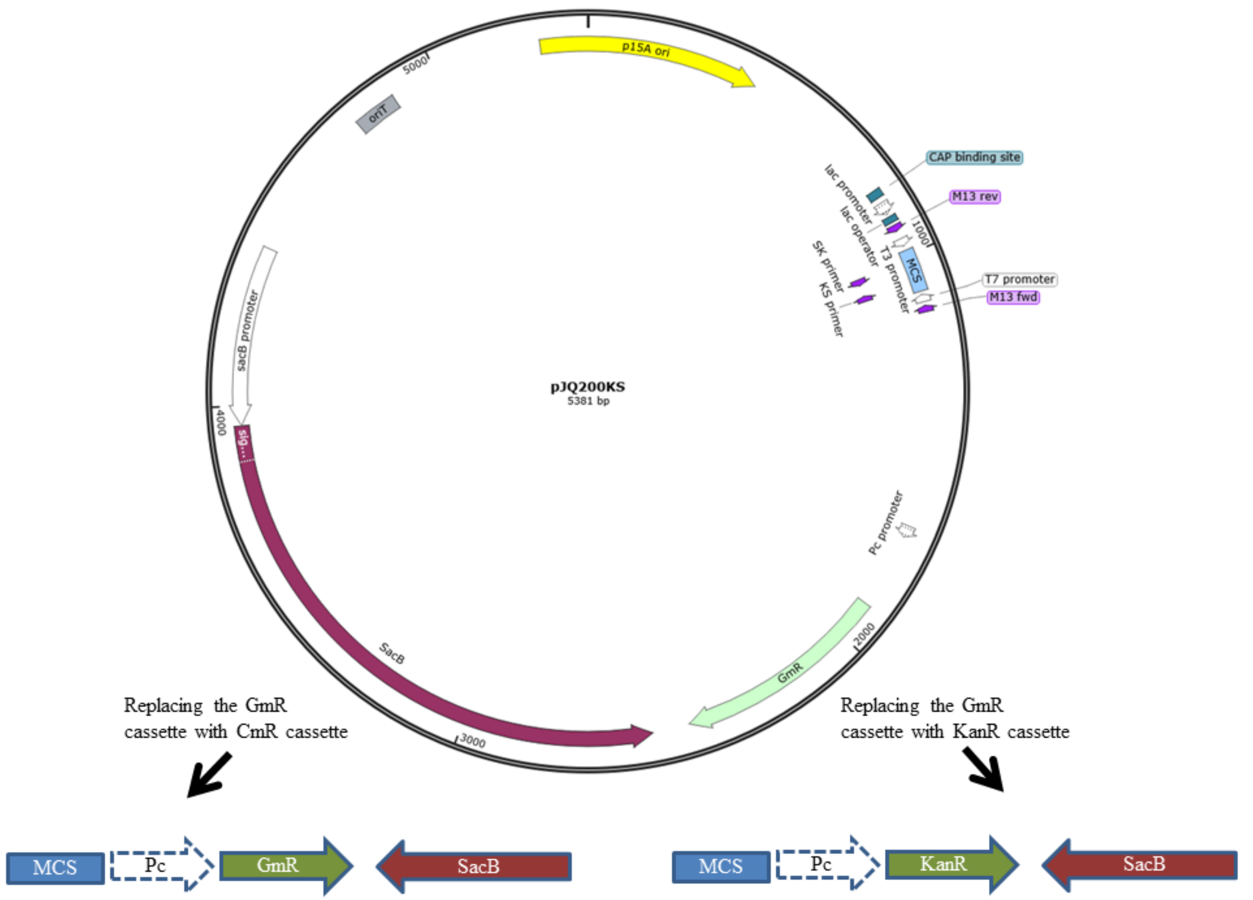


**Figure S3.** Creation of the two newly constructed pJQ200KS plasmids with gentamycin or kanamycin cassette **Gentamycin** (GmR) cassette of the pJQ200KS plasmid was replaced with either a kanamy.cin gene resistance cassette (KanR) or a chloramphenicol gene resistance cassette (CmR). MCS- Multiple Cloning Site. Pc-Pc- Pc promoter

**Figure S4.**

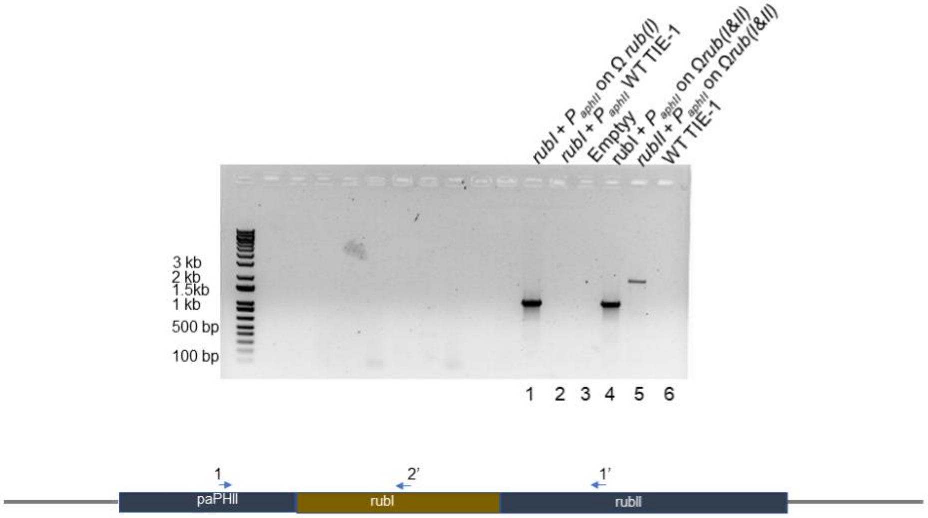


| |
|---|
| **Figure S4.** Checking the Ω*rub(I)* and Ω*rub(II)* engineered strains using PCR amplification spanning from the promoter region $P_{aphII}$ to the integrated either Rubisco form I (*rubI*)or form II (*rubII*). Expected band: Lane 1 ~800 bp, Lane 2=0 bp, Lane 3= empty, Lane 4 ~800 bp, Lane 5 ~1500 bp, Lan 6=0 bp. The primers set used for the PCR test are shown at the bottom of the figure and listed in Table S4. |

#### Supplementary Tables:

**Table S1.** List of antibiotics used in this study and their respective concentration.

| Antibiotic | Concentration (μg/mL) for *E. coli* | Concentration (μg/mL) for TIE-1 |
|---|---|---|
| Gentamycin | 20 | 200 |
| Chloramphenicol | 25 | 100 |
| Ampicillin | 100 | NA |
| Kanamycin | 50 | 50 |
| Tetracycline | 10 | NA |
| Sucrose | 100000 | 100000 |

**Table S2.** List of primers used to design plasmids used in this study.

| Primers used to design the plasmids used in this study | | Purpose |
|---|---|---|
| glmUSX_UP attB NcoI R | AAACCATGGtacgcgcccggggagcccaagggcacgccctggcacccGTCGCGAACTAGTAATGA | pWB083 |
| Histag AatII R | ATCGACGACGTCTAATGGTGATGGTGATGGTG | pWB081 |
| mCherry fd XbaI R | gcgcgcTCTAGACTCCAAAAAAAAAGGCTCCAAAAGGAGCCTTTAATTGTATCG ACGTCTAATGGTGATGGTG | pWB081 |
| attP paphII sacI F | ATATATGAGCTccccaactggggtaacctttgagttctctcagttggggCGCTAGCTTCACGCTG | pWB081 |
| mCherry NotI F | atatatGCGGCCGCAGGAGGatatacatATGGTGAGCAAGGGCG | pWB081 |
| paphII NotI R | ATATATGCGGCCGCGCGAAACGATCCTCA | pWB081 |
| PhiC31 NdeI F redesign | ATCGCCATATGgtggacacgtacgc | pWB084 |
| phi-C31 SacI R | atatatGAGCTCctacgccgctacgtct | pWB084 |
| LacIq NcoI F | ATGCGCCCATGGCTGATTGACACCATCGAATG | pWB084 |
| lacIq PciI F | GCGCGAACATGTCTGATTGACACCATCGAATG | pWB086 |
| φC31 NdeI F | ATATAGCATATGgtggacacgtacgcg | pWB088 |
| phi-C31 SacI R | atatatGAGCTCctacgccgctacgtct | pWB088 |
| KmR_PciI_AscI_F | agagACATGTaggaggGGCGCGCCatgattgaacaagatggattgcac | pWB091 |
| NdeI-KmR_RP | agagCATATGtcagaagaactcgtcaagaaggc | pWB091 |
| Cm from pBBRMCS1 AscI RBS p[illegible] | atcgatACATGTaggaggGGCGCGCCatggagaaaaaaatcactggatatacc | pWB092 |
| Cm from pBBRMCS-1 NdeI R | ATCGATCATATGttaatgaatcggccaacgcgcg | pWB092 |
| cbbL NdeI F | AACAGCATATGAACGAAGCAGTCACC | RuBisCO I |
| cbbLS fd XbaI R | atcgcgTCTAGAgTCCAAAAAAAAAGGCTCCAAAAGGAGCCTTTAATTGTATCTC | RuBisCO I |
| cbbM NdeI F | atcgacCATATGGACCAGTCGAAC | RuBisCO II |
| cbbM fd XbaI R | atcgTCTAGAgTCCAAAAAAAAAGGCTCCAAAAGGAGCCTTTAATTGTATCTTAC | RuBisCO II |

**Table S3.** List of primers used to design and check the mutants constructed in this study.

| Primer names | Sequence |
|---|---|
| Primers used to design and to check the *phaR* mutant | |
| *phaR* 1kb DN BamHI Fw 3 | tagttaGGATCCCAGCGAGGTTTGCTGCTTAG |
| *phaR* 1kb DN PstI Rev 4 | tcatcgCTGCAGATCGGGCCCGGTATCGCTCAC |
| *phaR* 1kb up BamHI Rev 2 | tcagtaGGATCCGTCACCCACACACACGGAGGA |
| *phaR* 1kb up SpeI Fw 1 | tacgtaACTAGTTCTTGCCGAGCTTCTCCGCCTGCTT |
| *phaR* gene Check Fw | ATGGCGAAATCAGACCAACCGAC |
| *phaR* gene Check Rv | CTATTCGTCCTTCTTCGGCTGT |
| *phaR* Check Fw | GTCAATCCGGCACACGCATCT |
| *phaR* Check Rev | GCAAAGCCGTCAATTAGCAAG |
| *phaR* 1kb Up check Fw | TCGTTGATTCCCGACGCAGAAC |
| *phaR* 1kb Up Check Rev | TCGAAGACATCCGGTTCTGACATG |
| *phaR* 1kb Dn Check Fw | CTTATCCTCCGTGTGTGTG |
| *phaR* 1kb Dn check Rev | CGGACTGGTGCTGAAGAACT |
| Primers used to design and to check the *phaZ* mutant | |
| *phaZ* 1Kb DN BamHI Rev 3 | ctacgtGGATCCTTACTACGCGTCGTCGTTAGCTG |
| *phaZ* 1Kb DN SpeI Rev 4 | tgagtACTAGTAGGTCGTTGCCGTAGGAGTCGA |
| *phaZ* 1Kb Up BamHI Rev 2 | tactctGGATCCCGTGTCGTGTCCCTCGTAAC |
| *phaZ* 1Kb Up SpeI Fw 1 | gtcatgACTAGTAGGTCGCATCTGGAAAGTTATCGC |
| *phaZ* gene Check Fw | CCCAACTGCTTCAACCTG |
| *phaZ* gene Check Rv | ATAGCCGCAGACCTCGTTCT |
| *phaZ* Check Fw | CGAATTCGTCCACGATGAG |
| *phaZ* Check Rev | TCGTCGACACCGAGAATG |
| *phaZ* check 1Kb Dn Fw | TGTGGACTGTCCGTGCTTAC |
| *phaZ* check 1Kb Dn Rev | AGTGGTTCATCTCGACCAGATG |
| *phaZ* check 1Kb Up Rev | ATGTACACAAGTCATCAGGATG |
| *phaZ* check 1Kb up Fw | AGGAGGTGTTTCTCGACGTC |

**Table S4.** Primers used to check the insertion of RuBisCO I and II in TIE-1 used in this study.

| Primer name | Sequence |
|---|---|
| $P_{aphII}$ Check Fw (1) | GTAGAAAGCCAGTCCGCAGAA |
| RuBisCO I Check Rev (2) | GGCGGGACATAGAAGTCGTA |
| RuBisCO II Check Rev (1') | TGTAGCCGATCACGAGGTC |
